# The 3.2Å resolution structure of human mTORC2

**DOI:** 10.1101/2020.04.10.029835

**Authors:** Alain Scaiola, Francesca Mangia, Stefan Imseng, Daniel Boehringer, Karolin Berneiser, Mitsugu Shimobayashi, Edward Stuttfeld, Michael N. Hall, Nenad Ban, Timm Maier

## Abstract

The protein kinase mammalian target of rapamycin (mTOR) is the central regulator of cell growth. Aberrant mTOR signaling is linked to cancer, diabetes and neurological disorders. mTOR exerts its functions in two distinct multiprotein complexes, mTORC1 and mTORC2. Here we report a 3.2 Å resolution cryo-EM reconstruction of mTORC2. It reveals entangled folds of the defining Rictor and the substrate-binding SIN1 subunits, identifies the C-terminal domain of Rictor as the source of the rapamycin insensitivity of mTORC2, and resolves mechanisms for mTORC2 regulation by complex destabilization. Two novel small molecule binding sites are visualized, an inositol hexakisphosphate (InsP6) pocket in mTOR and an mTORC2-specific nucleotide binding site in Rictor which also forms a zinc finger. Structural and biochemical analyses suggest that InsP6 and nucleotide binding do not control mTORC2 activity directly but rather have roles in folding or ternary interactions. These insights provide a firm basis for studying mTORC2 signaling and for developing mTORC2-specific inhibitors.

## Main Text

The serine/threonine kinase mTOR, a phosphatidylinositol 3-kinase-related kinase (PIKK) (*1-3*), controls cell growth by balancing anabolic and catabolic metabolism (*1, 4*). mTOR is found in two separate complexes: mTOR complex 1 (mTORC1) and mTORC2 (*1, 5, 6*). mTORC1 consists of mTOR, regulatory-associated protein of mTOR (Raptor), and mammalian homolog of protein lethal with sec thirteen protein 8 (mLST8) (*5, 7-9*). mTORC2 comprises mTOR, rapamycin-insensitive companion of mTOR (Rictor) (*10, 11*), stress-activated map kinase-interacting protein 1 (SIN1) (*12, 13*), and mLST8 (*10, 11*), and associates with the facultative subunit protein observed with Rictor-1/2 (Protor-1/2) (*14, 15*). mTORC2 is activated by insulin and phosphoinositide 3-kinase (PI3K) signaling (*6, 16*) and acts on cell survival and proliferation (*4*) by phosphorylating the AGC family kinases: Akt, PKC and SGK (*1, 4, 17-19*). mTORC2 also promotes tumorigenesis via upregulation of lipid biosynthesis (*20*).

mTOR inhibitors played a major role in the elucidation of mTOR signaling and are used in cancer treatment (*21*). The polyketide rapamycin specifically inhibits mTORC1 (*7, 8*) by forming a complex with the cellular protein FKBP12 that then binds the FKBP-rapamycin binding (FRB) domain in mTOR (Fig. 1) (*22*). ATP-like inhibitors target the ATP-binding site in the kinase catalytic domain of the mTORCs or the structurally related PI3K (*23*). Recently, mTORC2-selective inhibitors were identified, but their mechanism of action remains unknown (*24, 25*). Several intermediate resolution reconstructions of (m)TOR complexes (*26-30*) and high-resolution reconstructions of human mTORC1(*31*) have been reported, but no high-resolution information on mTORC2 is available. Of the mTORC2 accessory proteins, only the isolated pleckstrin homology (PH) and CRIM domains of SIN1 have been structurally characterized (*32-34*). For Rictor, fold- and secondary structure-based models have been proposed based on intermediate resolution cryo-electron microscopy (cryo-EM) reconstructions (*28-30*).

**Fig. 1.**
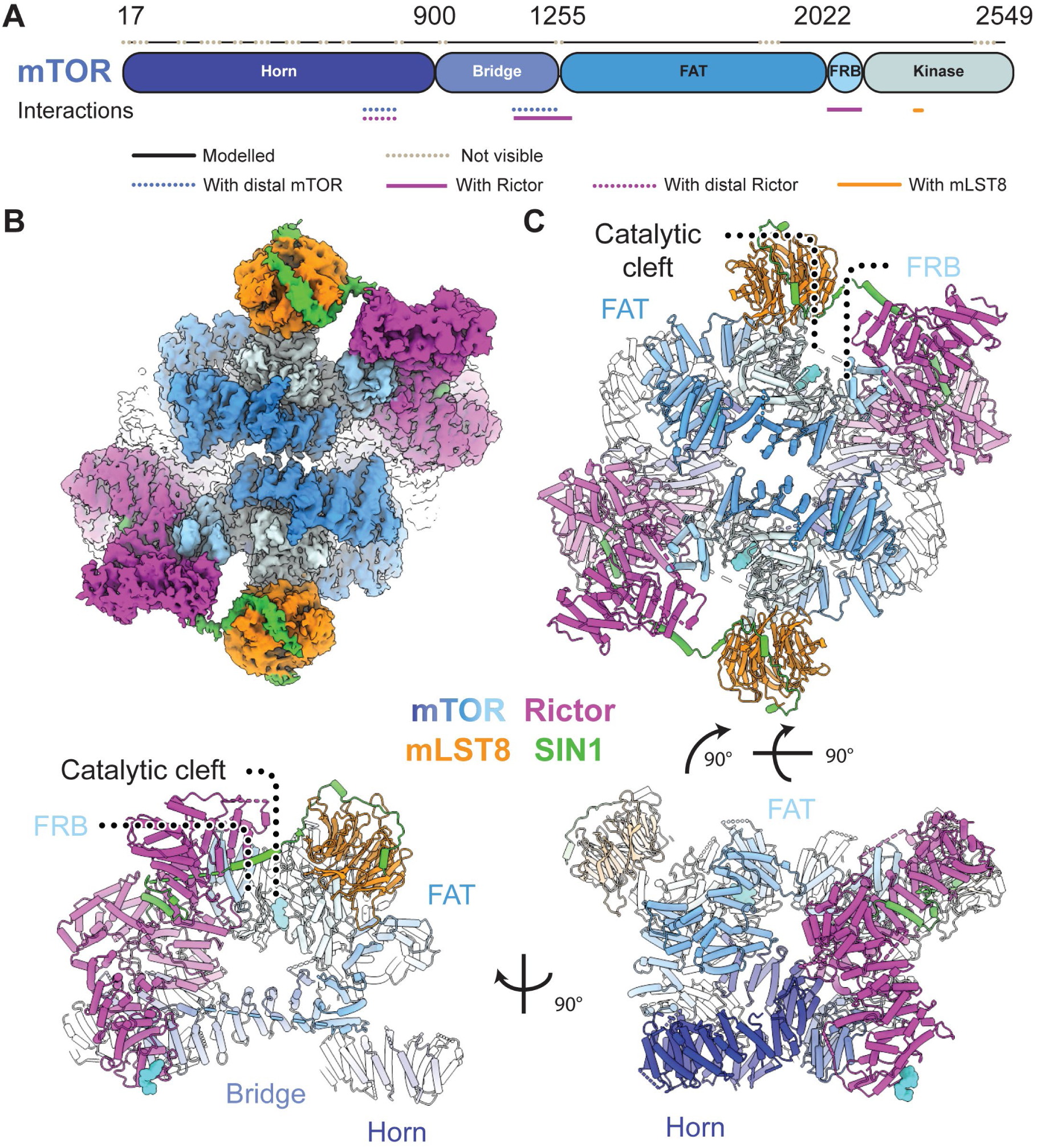
Structure of mTOR complex 2. (**A**) Sequence-level domain organization of mTOR. Modelled and unresolved regions are indicated as dotted lines. Interactions with other proteins in the complex are highlighted below the sequences. (**B**) Density of the overall cryo-EM reconstruction of mTORC2 colored according to protein subunits and mTOR domains as indicated. The top half is better resolved than the lower one, most likely due to conformational flexibility. (**C**) Cartoon representation of mTORC2 in three different orientations. The proteins Rictor (magenta) and SIN1(green) are unique to mTORC2, while mTOR (colored by domain) and mLST8 (orange) are common to both mTORC1 and mTORC2. Bound ligands are represented as cyan spheres.

To investigate the structure of mTORC2 and the mechanism of its regulation, we co-expressed recombinant components of human mTORC2 (mTOR, mLST8, Rictor and SIN1) in *Spodoptera frugiperda* cells. The assembled complex, purified using tag-directed antibody affinity followed by size exclusion chromatography, was analyzed by cryo-EM (Fig. 1B, and Figs. S1 and S2) in the presence of ATPγS and either the full-length substrate Akt1 (Fig. S3) or an Akt1 variant missing the PH domain (ΔPH-Akt1), or in the absence of Akt1 with and without ATPγS (Fig. S2). The sample prepared in the presence of ATPγS and ΔPH-Akt1 yielded the highest overall resolution of 3.2 Å (Density A in Fig. S2).

mTORC2 forms a rhomboid-shaped dimer (Fig. 1C) as observed in lower resolution mTORC2 reconstructions (*28-30*). The mTOR kinase forms the core of mTORC2 with mLST8 on the periphery, close to the active site cleft, similar to mTOR-mLST8 in mTORC1 (*26, 31*). In the overall reconstruction, as a consequence of EM refinement of a flexible molecule, one half of the dimer showed better local resolution (Fig. 1B and Figs. S4A-C and Movie S1). Therefore, focused refinement on a unique half of the assembly improved the resolution to 3.0 Å (Density C in Fig. S2), and these maps were used for structural modelling (Fig. S4D-F). Previous mTORC2 and yeast TORC2 reconstructions (*28-30*) revealed that the two mTOR FAT domains are in closer proximity to each other than observed in mTORC1 (*26, 31, 35*) and in the current structure, the distance between the mTOR FAT domains is further reduced (Fig. S5A). Irrespective of these structural differences between the two mTORCs, the catalytic site in mTORC2 closely resembles the catalytic site in mTORC1 without Rheb-mediated activation (*31*), suggesting that mTORC2 may be activated by a yet to be defined mechanism.

Previous studies of mTORC2 subunits Rictor and SIN1 or their yeast orthologs were not of sufficient resolution to allow de novo model building, resulting in ambiguous or inconsistent interpretations (*28, 30, 36*). Here we unambiguously model all structured regions of Rictor and the N-terminal region of SIN1 (Fig. 2A-C), whereas the middle and C-terminal part of SIN1 retain high flexibility and are not resolved. The fold of Rictor differs substantially from previous interpretations (*28*) (Fig. S5B-C). Rictor is composed of three interacting stacks of α-helical repeats, here referred to as the ARM domain (AD), the HEAT-like domain (HD), and the C-terminal domain (CD) (Fig. 2A-C and Fig. S6A). The N-terminal AD (residues 26-487) forms a large superhelical arrangement of nine ARM repeats (Fig. 2A-B) that structurally separates the HD and CD. The HD (residues 526-1007), interpreted as two separate domains in previous lower resolution studies (*28, 30*), is composed of ten HEAT-like repeats. In sequence space, the HD and CD of Rictor are separated by an extended stretch of residues (1008-1559) that are predicted to be disordered (*37*) and are not resolved in our reconstruction. We refer to this region as the phosphorylation-site region (PR) because it contains most of Rictor’s phosphorylation sites (*38*). The two ends of the PR are anchored by a two-stranded β-sheet at the top of the HD, which is thus termed the PR anchor (Fig. 2B-C and Fig. S6A). From here, a partially flexible linker wraps around the AD and the mTOR FRB domain extending toward the CD (Fig. 2B and Fig. S6C).

**Fig. 2.**
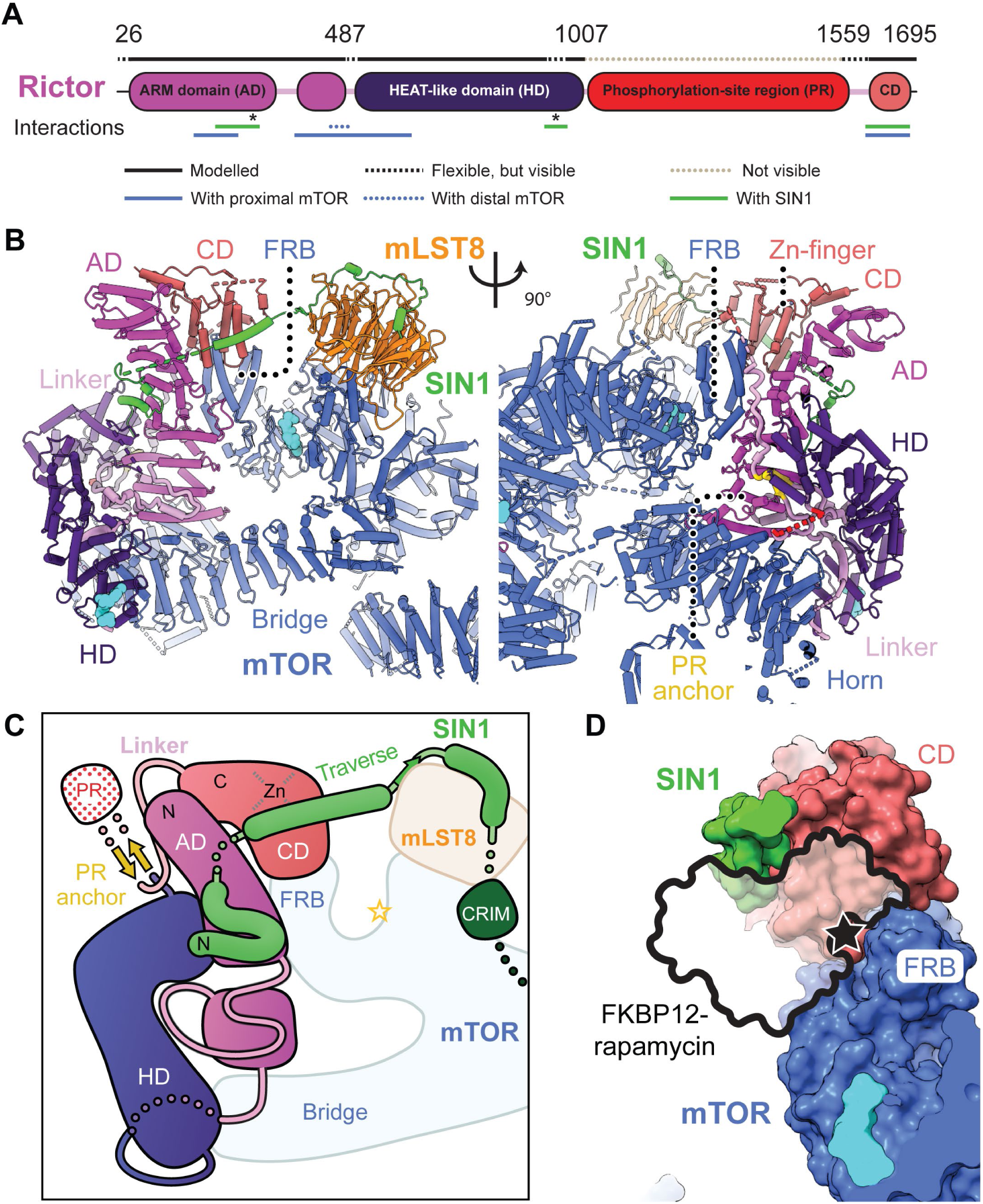
The architecture of Rictor. (**A**) Sequence-level domain organization of Rictor. Flexible and unresolved regions are indicated as dotted lines. Interactions with other proteins in the complex are highlighted below the sequences. Asterisks indicate residues interacting with the N-terminal region of SIN1. (**B**) Two views of Rictor, colored by domains. The structured part of Rictor forms three domains: an N-terminal Armadillo repeat domain (AD, magenta), a HEAT-like repeat domain (HD, dark magenta), and a C-terminal domain (CD, light red), the phosphorylation site region (PR) remains disordered. The sequences flanking the non-resolved PR are highlighted in red, the PR anchor is colored in gold. Bound ligands are shown as cyan spheres. (**C**) Schematic representation of Rictor and SIN1 domain topology. (**D**) The Rictor CD occupies the FRB domain and sterically blocks FKBP-rapamycin binding(*26*).

The structured parts of the CD form a four-helix bundle and a zinc finger, with bound Zn2+, in the vicinity of the Rictor N-terminus (Fig. 2A and Fig. S6B). Residues coordinating the zinc ion are highly conserved in metazoan Rictor (Fig. S6F). In earlier work, this domain had been interpreted as representing the SIN1 domain (*28*). The complete CD is absent in sequences of fungal Rictor orthologs, but other large extensions in yeast Rictor and SIN1 sequences may occupy the equivalent location in yeast TORC2, as observed in an intermediate resolution reconstruction of budding yeast TORC2(*29*) (Fig. S6D-E). Increased levels of Zn2+ have been reported to stimulate Akt S473 phosphorylation in cells (*39-41*), but no direct involvement of mTORC2 activation has been demonstrated.

Contacts between Rictor and mTOR are made by the Rictor AD, which sits between the proximal mTOR central HEAT domain and the N-terminal HEAT repeat domain, of the distal mTOR subunit (Fig. 2B). Due to its positioning on top of the mTOR FRB domain, the CD of Rictor blocks binding of FKBP12-rapamycin to mTORC2, thereby explaining mTORC2’s insensitivity to rapamycin (*5, 10, 11, 36*) (Fig. 2D).

The SIN1 subunit of mTORC2 exhibits an unexpected structural organization. The N-terminal region (residues 2-137), contrary to earlier interpretations, does not form an independently folding domain but interacts tightly with Rictor and mLST8 in an extended conformation (Fig. 2A-C and 3A-E). The CRIM, RBD and PH domains of SIN1, however, remain flexibly disposed. The N-terminus of SIN1 is inserted into a deep cleft at the interface of the AD and HD of Rictor. The N-terminal Ala2 with a structurally resolved acetylated N-terminus, and Phe3 of SIN1 are buried in a hydrophobic pocket of Rictor (Fig. 3C,D and Fig. S7A). The anchored N-terminal region of SIN1 forms two short helices (residues 6 to 33) inserted into grooves on the surface of the Rictor AD (Fig. 3D) and then continues with a flexible sequence segment toward the Rictor CD (Figs. 2B-C and 3C and Fig. S7B). Protruding from the Rictor CD, SIN1 forms a helical segment, referred to as the “traverse”, that spans the distance to mLST8 across the mTORC2 kinase cleft (Fig. 3C and Fig. S7B-C). The next region of SIN1 interacts with the fourth strand of the second blade of the mLST8 propeller by β-strand complementation, leading to displacement of an mLST8 loop relative to the structure of mLST8 in mTORC1 (Fig. 3C,E and Fig. S7D). SIN1 then follows the surface of the mLST8 propeller, finally forming an α-helix anchored between the first and seventh blades of mLST8.

**Fig. 3.**
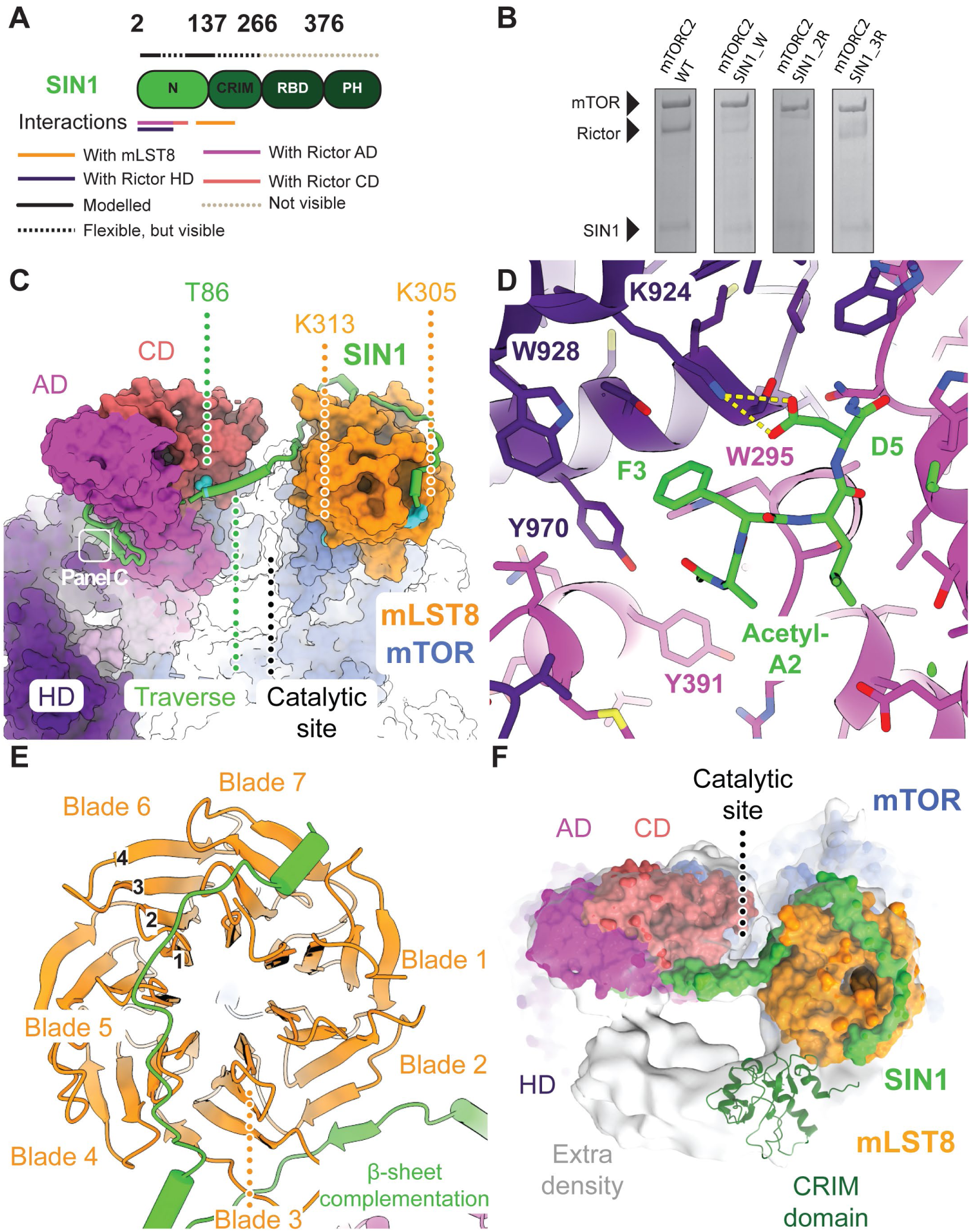
The SIN1 N-terminal region is an integral component of mTORC2. (**A**) Sequence-level domain organization of SIN1. Flexible and unresolved regions are shown above each domain representation as dotted lines in two colors as indicated. Interactions with other proteins in the complex are indicated below the domain representation. (**B**) Extension of the processed SIN1 N-terminus disrupts assembly of Rictor and SIN1 with mTOR/mLST8 into mTORC2. SDS-polyacrylamide gel of a FLAG bead pulldown from lysates of insect cells expressing mTORC2 comprising SIN1 variants. Levels of Rictor are drastically reduced in the mTOR-based pulldown for mTORC2 carrying variants of SIN1 N-terminally extended by a tryptophan (mTORC2 SIN1_W), two consecutive arginines (mTORC2 SIN1_2R) and three consecutive arginines (mTORC2 SIN1_3R) (**C**) Surface representation of mTORC2. SIN1 (shown as green cartoon) interacts via two N-terminal helices with Rictor, winds around Rictor, traverses the catalytic site cleft and winds around mLST8. The field of view of subpanel C is indicated. (**D**) Close-up view of the SIN1 N-terminal residues, which are deeply inserted between Rictor AD and HD. Acetylated Ala2 and Phe3 are bound in a hydrophobic pocket, while Asp5 interacts via salt bridges (yellow dashes). (**E**) Top view of mLST8 β-propeller (orange) and the interaction regions with SIN1 (green). The nomenclature for WD40 β-propeller repeats is indicated. (**F**) Top view of the catalytic site with the structure shown as surface together with the density of a subclass (light grey). The lower resolution extra density is consistent with a placement of the SIN1 CRIM domain, here shown in dark green (PDB: 2RVK). Unassigned extra density protrudes from the CRIM domain to the mTOR active site and Rictor.

SIN1 integrates into the Rictor fold and connects Rictor with mLST8, suggesting a direct role in stabilizing mTORC2. To test the relevance of the anchoring of the N-terminus of SIN1 on Rictor, we extended the N-terminus of SIN1. Insertion of residues impairs critical interactions observed for the acetylated N-terminus of SIN1 and prevents Rictor integration into mTORC2, as observed in Baculovirus-mediated expression of mTOR components followed by pull-down assays (Fig. 3B and Fig. S8). Therefore, SIN1 acts as an integral part of the Rictor structure that critically stabilizes interdomain interactions, explaining the difficulties observed in purifying isolated Rictor (*28*).

These observations are also consistent with the locations of post-translational modifications or mutations that affect mTORC2 activity. SIN1 phosphorylation at Thr86 and Thr398 has been reported to reduce mTORC2 integrity and kinase activity toward Akt Ser473 (*42*). Thr86 in SIN1, which is a target for phosphorylation by S6 kinase (*42*), is bound to a negatively charged pocket of the Rictor CD (Fig. 3C and Fig. S7C). Phosphorylation of Thr86 would lead to repulsion from this pocket, destabilizing the interaction between Rictor and mTOR-mLST8 and presumably the entire mTORC2 assembly, in agreement with earlier in vivo and in vitro observations (*42*). The importance of SIN1 in connecting Rictor to mLST8 and, therefore also indirectly to mTOR, is also consistent with the requirement of mLST8 for mTORC2 integrity (*43, 44*).

A poorly resolved density linked to the SIN1 helix anchored to mLST8 is observed in all reconstructions. In previous structural studies of yeast TORC2, a similar region of density was associated with the CRIM domain of Avo1, the yeast SIN1 ortholog (*29, 36*). Most likely, it represents the mobile substrate-binding CRIM domain that directly follows the helix in the SIN1 sequence and has a matching shape based on the solution structure of the S. pombe SIN1 CRIM domain (*33, 34*) (Fig. 3C-F and Fig. S9A,C). The positions of the SIN1 RBD and PH domains remain unresolved. In the dataset collected for samples with added full-length Akt1 (Dataset 2 in Fig. S2), we observed additional low-resolution density (Fig. 3F and Fig. S9B-C) between the hypothetic CRIM domain and Rictor AD and CD in the vicinity of the mTOR active site. This density, not of sufficient resolution to assign specific interactions, may represent parts of bound Akt1 or SIN1 domains (Fig. S9C).

A proposed regulatory mechanism for mTORC2 involves ubiquitylation of mLST8 on Lys305 and Lys313 (*45*). Loss of ubiquitylation by K305R and/or K313R mutation, or truncation of mLST8 at Tyr297, leads to mTORC2 hyperactivation and increased Akt phosphorylation (*45*). Indeed, mLST8 Lys305 is proximal to the SIN1 helix anchoring the CRIM domain. Ubiquitylation of Lys305 would prevent association of the SIN1 helix, leading to dislocation of the SIN1 CRIM domain required for substrate recruitment (Figs. 3C and 4C). Ubiquitylation of Lys313, which is found on the lower face of mLST8 (Figs. 3C and 4C), presumably also interferes with positioning of the CRIM domain (Fig. S9).

We observed two novel, small molecule binding sites outside the mTOR catalytic site, which is itself occupied by ATPγS. The first (A-site) (Fig. 4A and Fig. S10B) is located in the HD of Rictor and is thus specific to mTORC2. The second (I-site) (Fig. 4B and Fig. S10C) is located in the FAT domain of mTOR and is thus common to mTORC1 and mTORC2.

**Fig. 4.**
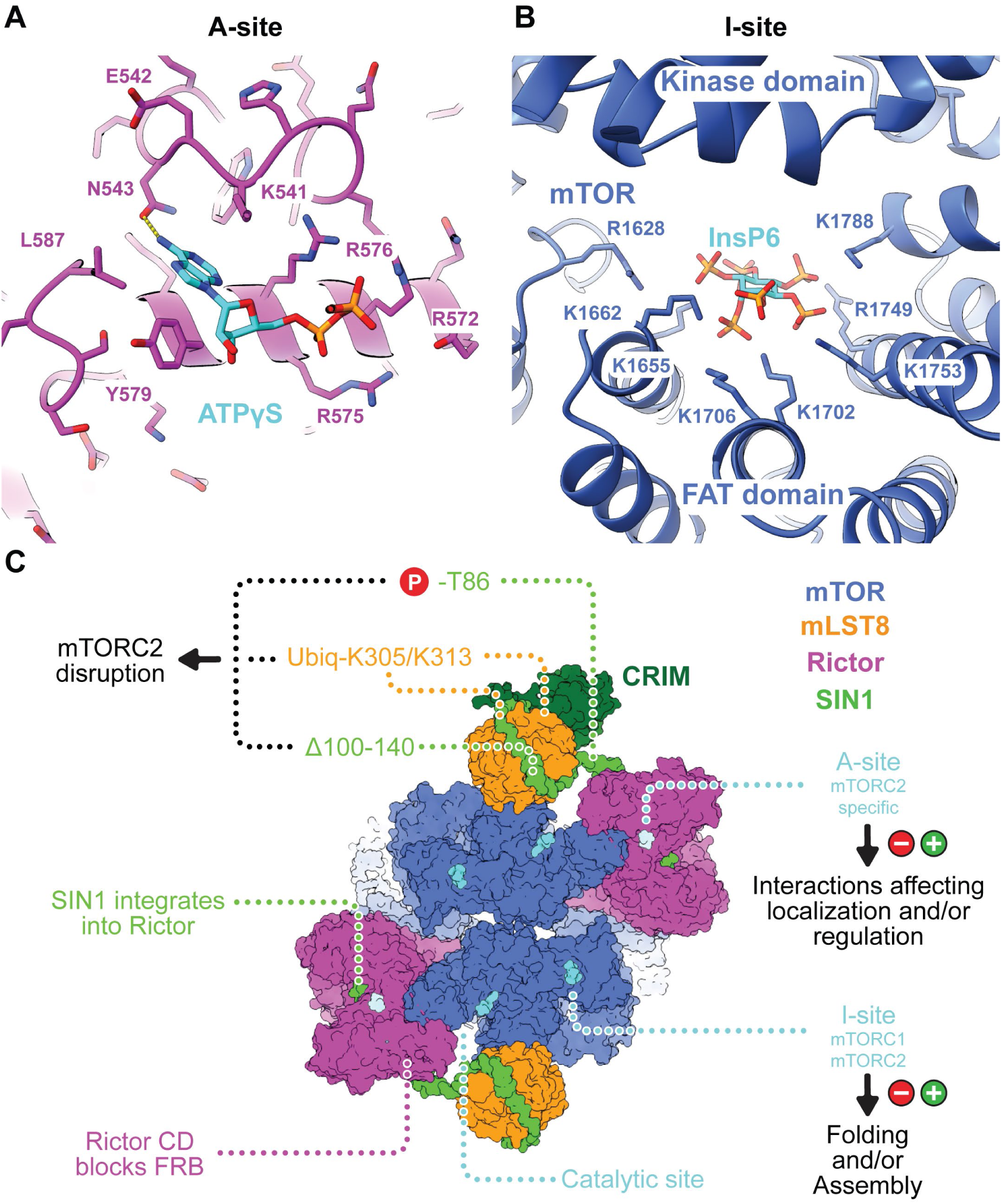
Small molecule binding sites of mTORC2 outside the active site region. (**A**). Close-up view of the A-site on the periphery of the Rictor HD with bound ATPγS. A hydrogen bond between ATPγS and Asn543is shown as dashed yellow lines. (**B**) Close-up view of the I-site in the FAT domain of mTOR. InsP6 is surrounded by a cluster of positively charged amino acids. It only directly interacts with residues of the FAT domain. (**C**) Overview of mTORC2 architecture and ligand interaction sites. Each half of the dimeric mTORC2 has three small molecule binding sites. The kinase active site and the A-site, which is located in the peripheral region of Rictor, bind to ATP (or ATP analogues). The I-site in the middle of the FAT domain of mTOR binds InsP6. The indicated modifications on SIN1 and mLST8 affect mTORC2 assembly. Extra-density region following the CRIM domain is indicated as a grey outline.

The density of the small molecule in the A-site matched that of an ATP molecule and was confirmed to be ATP (or ATPγS) through a comparison of cryoEM reconstructions of mTORC2 with and without ATPγS added at a near physiological concentration of 2mM (Datasets 1 and 4, Fig. S2 and S10A). The A-site does not resemble any known ATP binding site. Positively charged amino acids (Lys541, Arg575, Arg576, Arg572) of the A-site are conserved in Rictor orthologs from yeast to human (Figs. S6E and S11). Other residues are not conserved, hinting at the possibility for interactions with alternative negatively charged ligands. The A-site is located approximately 100 Å from the mTOR catalytic site. Ligand binding to the A-site caused neither long-range allosteric change affecting the kinase site nor local structural perturbations (Fig. S12).

To investigate the effect of ligand binding to the A-site, we generated a series of Rictor variants with a mutated A-site (Table S1). Variants with three or four mutated residues (A3 and A4) assembled into mTORC2 (Fig. S13B) while variant A5 was defective in assembly (Fig. S13B-D). Cryo-EM reconstructions of variants A3 and A4 in the presence of ATPyS (Fig. S12I-J,K-L) confirmed that the chosen mutations abolish ligand binding under near physiological conditions (Figs. S10A and S12J,L). Purified mTORC2 containing Rictor variants A3 or A4 exhibited thermal stability and kinase activity, in an Akt1 in vitro phosphorylation assay, comparable to wild-type mTORC2. (Figs. S14B and S15A,B). Complementation of a Rictor knockout (KO) in HEK293T cells by transfected Rictor-WT, or Rictor variant A3 yielded comparable levels of AKT-S473 phosphorylation (Table S1 and Fig. S16). Altogether, the above analyses indicate that ligand binding to the A-site does not directly influence mTORC2 kinase activity, suggesting rather a role in the interaction with other, yet unidentified, partner proteins of mTORC2.

The I-site is formed entirely by the FAT domain of mTOR, where a large, positively charged, pocket is lined by six lysine and two arginine residues to bind an extended ligand (Fig. 4B and Fig. S10C). The I-site was still partially occupied in our reconstruction of mTORC2 prepared without addition of exogeneous ATPγS or other relevant ligands (Data Fig. S10A). The co-purified molecule was identified by map appearance and by ion mobility spectrometry-mass spectrometry (IMS-MS) as inositol hexakisphosphate (InsP6) (Figs. S17 and S18A-F). InsP6 binds in a region, which is incomplete in related PI3Ks (*46*), but generally conserved in members of the PIKK family of kinases (*47*). Indeed, InsP6 was previously reported to associate with DNA-PKcs (*48*). Recently, structure determination of the PIKK family pseudo-kinase SMG1 revealed InsP6 binding in a region corresponding to the I-site and led the authors to postulate a corresponding binding site in mTOR but involving both the kinase domain and FAT domain (*47*). InsP6 has previously been observed as a structural component of multi-subunit assemblies, including the splicesome (*49*) and proteasome activator complex (*50*), and helical repeat regions have been identified as InsP6 interaction sites (*51*).

To investigate the function of Ins6P interaction, we purified recombinant mTORC2 containing mTOR I-site mutations (Table S1). mTOR variants with two and three mutations, I2 and I3, yielded intact mTORC2 complexes (Fig. S13A), while a variant with five mutations, I5, failed to assemble into mTORC2 (Fig. S13A,C). mTORC2 containing mTOR variants I2 and I3 displayed normal kinase activity toward Akt1 in vitro (Fig. S14A). Notably, the mutations in I2 are equivalent to those reported previously to abolish completely the kinase activity of an N-terminally truncated ‘naked’ mTOR fragment toward a C-terminal peptide of Akt1 (*47*). A possible explanation for this apparent discrepancy is provided by a reduced stability of mTORC2 assembled using the I2 variant (but not the I3 variant) (Fig. S15A). This destabilizing effect might be more pronounced in an mTOR fragment than in the context of an assembled mTORC2 (Fig. S15A).

To investigate a possible role of InsP6 metabolism on mTORC2 activity in HEK293T cells, we knocked down (KD) and knocked out (KO) Inositol-pentakisphosphate 2-kinase (IPPK) and Multiple inositol polyphosphate phosphatase 1 (MINPP1), respectively. The former enzyme generates InsP6 whereas the latter degrades it (Fig. S19). These manipulations of InsP6 metabolizing enzymes did not alter mTORC2 kinase activity in non-stimulated cells or in cells stimulated with FCS and insulin (Fig. S19A-H). These biochemical results are consistent with the observed stable binding of InsP6 to mTORC2 and suggest a role of InsP6 in mTOR folding or mTOR complex assembly, rather than as an acute transient metabolic input signal to mTORC1 or mTORC2.

Here, we describe a bona fide structure of mTORC2. We visualized how SIN1 stabilizes and tethers Rictor to the mTOR-mLST8 core. SIN1 further uses mLST8 as a platform for positioning its substrate recruiting CRIM domain, revealing a new functional role for mLST8 and rationalizing the impact of SIN1 and mLST8 modifications on mTORC2 activity. We also provide the structural basis for how the Rictor CD determines mTORC2’s rapamycin insensitivity, by a mechanism different from those inferred from previous structural data (*28, 30*). We identified and functionally characterized two ligand binding sites in mTORC2. The I-site in mTOR is common to mTORC1 and 2, binds InsP6 and presumably functions in mTOR folding or assembly rather than acting as a sensor site for acute changes in cellular InsP6 concentration. The mTORC2 specific A-site of Rictor binds ATP. It doesn’t affect mTORC2 activity by allostery but may be involved in linking partner protein interactions to cellular nucleotide triphosphate concentrations. Altogether, the data presented here provide a firm basis for further analysis of the function of mTORC2 and its interplay with partner proteins for controlling subcellular localization (*52*) and regulation of activity (*1, 4, 10, 20*). Interaction sites of Rictor and mLST8 with SIN1 provide an opportunity for the development of inhibitors specific for mTORC2.

## Supporting information

Material and Methods and all supplementary materials

## Acknowledgments

We thank T. Sharpe at the Biophysics facility and A. Schmidt at the Proteomics Core Facility of Biozentrum and the sciCORE scientific computing facility, all of University of Basel. We thank M. Leibundgut for advice with model building, A. Jomaa and S. Mattei for advice on cryoEM data processing, the ETH scientific center for optical and electron microscopy (ScopeM), and in particular M. Peterek and P. Tittmann for technical support. We are indebted to E. Laczko and J. Hu of the Functional Genomics Center Zürich for the help with the mass spectrometry. We thank Iva Lučić and Thomas Leonard (Max F. Perutz Laboratories, Vienna) for providing (Delta-PH) Akt1 protein.;

## Author contributions

AS designed the experiments, prepared the sample for cryoEM, carried data processing and structure modelling. AS and DB performed data collection. FM designed the experiments, cloned Akt1, mTORC2 mutants and Rictor mutants, expressed and purified proteins, performed the activity assays and the nanoDSF measurements. ES established the mTORC2 purification procedure. SI cloned mTORC2, contributed to data analysis and manuscript preparation. MS performed the in-cell analysis of mTORC2 activity. KB and MS performed the KO/KD of MINPP1 and IPPK. AS, FM, DB, SI, NB, MNH and TM participated in the writing of the manuscript.;

## Data and materials availability

The high resolution cryo-EM map of the half (Density C) and full-mTORC2 (Density A) has been deposited in the Electron Microscopy Data Bank as EMD-XXX and EMD-YYYY respectively, while the corresponding model are in the Protein Data Bank as PDB ID WWW and ZZZ. Additionally, the density of mTORC2 in absence of ATPγS (Density F), as well as the densities showing extra density (Density G and H) were deposited in the Electron Microscopy Data Bank as EMD-AAAA, EMD-BBBB, and EMD-CCCC respectively.

## Supplementary Materials

Materials and Methods

Figures S1-S20

Tables S1-S2

Movies S1-S4

External Databases S1-S7

