## Supplementary material for "The 3.2Å resolution structure of human mTORC2": Material and Methods and all supplementary materials

### Supplementary Materials for
### The 3.2Å resolution structure of human mTORC2

Alain Scaiola[2†], Francesca Mangia[1†], Stefan Imseng[1], Daniel Boehringer[3], Karolin Berneiser[1], Mitsugu Shimobayashi[1], Edward Stuttfeld[1], Michael N. Hall[1], Nenad Ban[2*], & Timm Maier[1*].

[†] These authors contributed equally to this work

[*] Correspondence should be addressed to N.B. or T.M.

**This PDF file includes:**

Materials and Methods

Supplementary Text

Figs. S1 to S20

Tables S1 to S2

Captions for Movies S1

**Other Supplementary Materials for this manuscript include the following:**

Movies S1-S4

### Materials and Methods

Protein expression and purification

Insect cell vectors from the ‘MultiBac’ Baculovirus expression system(*53*) (Geneva Biotech, Geneva, Switzerland) have been used to clone internally FLAG-tagged pAceBAC-mTOR (FLAG after Asp258), pIDK-Rictor, pIDC-mLST8, and pAceBAC1-SIN1 using Gateway Cloning (Thermo Fisher Scientific, US). Rictor was originally amplified from myc-Rictor, which was a gift from David Sabatini(*11*) (Addgene plasmid #11367). Site-directed mutagenesis was used to generate mTORC2 A- and I- site variants. The following set of A-site mutants with pIDK-Rictor as template was created: Rictor_R572E_R575E_R576E (A3), Rictor_R572E_R575E_R576E_Y579A (A4), Rictor_R572E_R575E_R576E_Y579A_L587W (A5). The following I-site mutants with FLAG-tagged pAceBAC-mTOR were generated: mTOR_K1753E_K1788E (I2), mTOR_R1628E_K1655E_K1662E (I3) and mTOR_R1628E_K1655E_K1662E_K1706E_K1735E (I5). Wild-type Rictor and mutants A3, and A5 were subcloned into a gentamycin resistant-mammalian expression vector under control of a CMV promoter. SIN1 N-terminal variants were generated by inserting a tryptophan (SIN1_W), two consecutive arginines (SIN1_2R) or three consecutive arginines (SIN1_3R) using site-directed mutagenesis and pAceBAC1-SIN1 as template. Plasmids encoding FLAG-tagged mTOR, Rictor and mLST8 were fused to a ‘MultiBac’ expression plasmid using Cre-recombinase (New England Biolabs, Ipswich, USA) and transposed into a bacmid for baculovirus production. Baculovirus encoding untagged SIN1 was produced separately.

Sf21 insect cells (Expression Systems) were grown in HyClone insect cell media (GE Life Sciences) and Baculovirus was generated according to the Fitzgerald et al., 2006(*53*). For the expression of recombinant human WT mTORC2, A- and I- site mTORC2 mutants and mTORC2 carrying SIN1 N-terminal variants, Sf21 cells were infected at a cell density of 1 Mio/ml. Cells were coinfected with 1:100 (v/v) ratio of two undiluted supernatants from cells previously infected with baculovirus encoding FLAG-mTOR, Rictor and mLST8, or infected with baculovirus encoding untagged SIN1, respectively. WT mTORC2, A- site mutants A3, A4 and A5 and I- site mutants I2, I3 and I5 were purified as follows: insect cells were harvested 72 hours post infection by centrifugation at 800 x g for 25 minutes and stored at −80 °C until further use. Cell pellets were lysed in 50 mM bicine pH 8.5, 200 mM NaCl, 2 mM $MgCl_2$ by sonication and the lysate cleared by ultracentrifugation. Soluble protein was incubated with 10 ml anti-DYKDDDDK agarose beads (Genscript, Piscataway, USA) for 1 hr at 4°C. The beads were transferred to a 50 ml gravity flow column (BioRad) and washed four times with 200 ml of wash buffer containing 50 mM bicine pH 8.5, 200 mM NaCl, 2 mM EDTA. Protein was eluted by incubating beads for 30 min with 10 ml wash buffer supplemented with 0.6 mg/ml synthetic DYKDDDDK peptide (Genscript, Piscataway, USA). The eluate was combined with three additional elution steps using 0.1 mg/ml synthetic DYKDDDDK peptide and five minutes incubation time. The eluted protein was concentrated using a 100,000 Da molecular mass cut off centrifugal concentrator (Amicon) of regenerated cellulose membrane and purified by size-exclusion chromatography on a custom-made Superose 6 Increase 10/600 GL gel filtration column equilibrated with 10 mM bicine pH 8.5, 150 mM NaCl, 0.5 mM EDTA, 2 mM TCEP. Purified WT mTORC2 was concentrated in gel filtration buffer to a final concentration of 3-3.5 mg/ml determined by A280 absorption using a NanoDrop 2000; Thermo Scientific. Sample was supplemented with 5% (v/v) glycerol and stored at -80 °C for later cryo-EM use. Purified mTORC2 variants with A- and I-site mutants were concentrated in gel filtration buffer to a final concentration of 0.4-2 mg/ml as determined by absorption at 280nm wavelength using a NanoDrop 2000 (Thermo Scientific). The resulting samples were supplemented with 5% (v/v) glycerol and stored at -80 °C for later use.

The coding sequence for Akt1(*54*), was cloned into a pAceBAC1 expression vector (Geneva Biotech, Geneva, Switzerland) with an N-terminal His10–Myc–FLAG tag by Gateway cloning. Baculovirus was produced as described for mTORC2. Akt1 was purified with anti-DYKDDDDK agarose beads as described for mTORC2. The eluted protein was concentrated using a 10,000 Da molecular mass cut off centrifugal concentrator (Amicon) of regenerated cellulose membrane and further purified by size-exclusion chromatography with a Superdex 75 Increase column equilibrated with 10 mM bicine pH 8.5, 150 mM NaCl, 0.5 mM EDTA, 2 mM TCEP. Purified Akt1 was concentrated in gel filtration buffer, supplemented with 5% (v/v) glycerol and stored at -80 °C for further experiments. Dephosphorylated Akt1 was obtained after overnight incubation of 4.5 mg of protein with 6 µg of λ-protein phosphatase (New England Biolabs) in presence of PMP buffer (New England Biolabs) and 1mM $MnCl_2$ prior to size exclusion chromatography. Successful Akt1 dephosphorylation was confirmed by Western Blot with antibodies against phospho-Akt-Ser473 (#4060 Cell Signaling Technologies, Beverly, USA) and phospho-Akt-Thr450 (#9267 Cell Signaling Technologies, Beverly, USA). Human (Delta-PH)-Akt1 protein (residues 144-480, mono-phosphorylated on T450), as described in Lucic et al.(*55*) (therein referred to as Akt1KD), was provided by T. Leonard (Max-Perutz Labs, Vienna).

<u>Expression and assembly analysis via immunoprecipitation</u>

A-site mutants A3, A4, A5, and I- site mutants I2, I3, I5 and mTORC2 carrying SIN1 N-terminal variants extended by a tryptophan (SIN1_W), two consecutive arginines (SIN1_2R) and three consecutive arginines (SIN1_3R) inserted between the processed Met1 and Ala2, were immunoprecipitated in small-scale using FLAG beads. Five grams wet weight of pellets from insect cells expressing A- and I- site mutants and SIN1 N-terminal variants were lysed in 50 mM bicine pH 8.5, 200 mM NaCl, 2 mM $MgCl_2$ using a Dounce homogenizer. The lysate was cleared by ultracentrifugation for 45 minutes at 35,000 x g. Soluble protein was incubated with 125 µl of anti- DYKDDDDK agarose beads (Genscript, Piscataway, USA) for 1 h at 4°C. The beads were transferred to a 5 ml gravity flow column (Pierce Centrifuge Columns, Thermo Scientific) and washed with 50 ml of buffer containing 50 mM bicine pH 8.5, 200 mM NaCl, 2 mM EDTA. Protein was eluted by 30 minutes incubation of the beads with 400 µl wash buffer supplemented with 0.6 mg/ml synthetic DYKDDDDK peptide Genscript, Piscataway, USA). Total lysate, soluble supernatant after ultracentrifugation, flowthrough from FLAG column, buffer wash and elution fraction were loaded onto a 4-15% SDS polyacrylamide gel (Bio-Rad Laboratories). Additionally, total lysate, supernatant after ultracentrifugation and elution fraction of mTORC2 WT, SIN1 N-terminal variants and mutants A5 and I5 were analysed by immunoblotting using antibodies against mTOR (#2972; Cell Signaling Technologies, Beverly, USA), SIN1 (Bethyl A300-910A), Rictor (Bethyl A300-458A) and actin (MAB1501; Merck Millipore).

Assay for mTORC2 kinase activity

mTORC2 kinase activity assays were conducted in 100 mM HEPES pH 7.4, 1 mM EGTA, 1 mM TCEP, 0.0025% Tween-20, 10 mM $MnCl_2$ using dephosphorylated Akt1 as a substrate. In a 60 µl reaction volume, 0.05 µM of either WT or A- and I- sites mutant mTORC2 were mixed with 1 µM Akt1 and, where indicated, either DMSO or 25 µM Torin1. The mixture was preincubated for five minutes at room temperature and the reaction was initiated by the addition of 10 µM ATP. After 20 minutes at 37°C the reaction was terminated by the addition of 60 µl 2x Laemmli sample buffer. The reactions were analysed by western blotting using primary antibodies against phospho-Akt-Ser473 (#4060; Cell Signaling Technologies, Beverly, USA), phospho-Akt-Thr450 (#9267; Cell Signaling Technologies, Beverly, USA), Akt (#4685), and mTOR (#2972; Cell Signaling Technologies, Beverly, USA), anti-FLAG antibodies (Sigma-Aldrich F1804), SIN1 (Bethyl A300-910A), Rictor (Bethyl A300-458A) at a dilution of 1:1000. A goat anti-rabbit HRP-labeled antibody (ab6721; Abcam, Cambridge, UK) was used as the secondary antibody at a dilution of 1:3000. Signals were detected using the Enhanced Chemiluminescence (ECL) kit SuperSignal™ West Femto Maximum Sensitivity Substrate (Thermo Scientific). Images were acquired using a Fusion FX (Vilber) imaging system.

Thermal stability assay

Thermal unfolding was monitored by Differential Scanning Fluorimetry (DSF) based on internal tryptophane fluorescence on a Prometheus NT.48 instrument (NanoTemper Technologies). Purified WT mTORC2 or mTORC2 containing mutations in A- or I- site were diluted to 0.1 mg/mL in 10 mM bicine pH8.5, 150 mM NaCl, 0.5 mM EDTA, 2 mM TCEP. High precision capillaries (NanoTemper Technologies) were filled with 10 µL sample and placed on the sample holder. A temperature gradient of 0.1°C/min from 22 to 65 °C was applied and fluorescence intensity at 330 and 350 nm was recorded. A plot of the ratio of fluorescence intensities at those wavelengths (F350/F330) was generated using a Python script. The experiment was repeated two times with five replicates per sample run each time. Melting points were calculated using PR.ThermControl software version 2.1.2. Data were analyzed using GraphPad Prism version 8.0.0 (GraphPad Software, San Diego, California USA) to generate the mean and SD of the melting points. One outlier, likely resulting from capillary handling, for sample A4 was excluded from data analysis.

In cell analysis of mTORC2 activity for A-site mutants

HEK293T cells were cultured and maintained in DMEM high glucose with 10% FCS, 4 mM glutamine, 1 mM sodium pyruvate, 1x penicillin/streptomycin. RICTOR knockout cells were generated as described in Bossler et al., 2019(*56*). 4 µg of plasmids harboring RICTOR-WT, RICTOR-A_3, and RICTOR-A_5 were transfected with JetPRIME (Polyplus). 24 hours after transfection, cells were starved for serum for overnight and stimulated with 10% FCS and 100 nM insulin for 15 min. Total cell lysates were prepared in lysis buffer containing 100 mM Tris-HCl pH7.5, 2 mM EDTA, 2 mM EGTA, 150 mM NaCl, 1% Triton X-100, complete inhibitor cocktail (Roche) and PhosSTOP (Roche). Protein concentration was determined by a Bradford assay, and equal amounts of protein were separated by SDS-PAGE, and transferred onto nitrocellulose membranes (GE Healthcare). Antibodies used were as follows: AKT (Cat#2920, Cell Signaling Technology), AKT-pS473 (Cat#4060, Cell Signaling Technology), RICTOR (Cat#2040, Cell Signaling Technology), ACTIN (Cat#MAB1501, Millipore), IRDye 800CW goat anti-rabbit IgG (Cat#926-32211, LI-COR), and IRDye 680RD goat anti-mouse IgG (Cat#926-68070). Signals were detected by LI-COR Fc (LI-COR Biosciences).

In cell analysis of the dependence of mTORC2 activity on IPPK and MINPP1

HEK293T cells were cultured and maintained in DMEM high glucose with 10% FCS, 4 mM glutamine, 1 mM sodium pyruvate, 1x penicillin/streptomycin. For knockdown of IPPK and MINPP1, 0.1 x 106 cells/well were seeded in a 6-well plate and transfected with 100 nM siRNA using the jetPRIME (Polyplus) system. After 32h, cells were washed twice with PBS (-/-) and starved for serum for 16 hours. 48 hours post-transfection, cells were incubated at 37 °C with PBS (+/+) for 10 min followed by stimulation with 10 % FCS and 100 nM insulin for 15 min at 37 °C. Cells were washed with ice-cold PBS (-/-) and harvested for SDS-PAGE or RNA isolation for qPCR analysis. Knockout experiments were conducted as described above, using generated KO cells instead of transfection with siRNA. Total cell lysates were prepared in M-PER lysis buffer (ThermoFisher) containing complete inhibitor cocktail (Roche) and PhosSTOP (Roche), and protein concentrations determined by Bradford assay. Equal amounts of protein were separated by SDS-PAGE, transferred onto nitrocellulose membranes (GE Healthcare), and signals were detected by LI-COR Fc (LI-COR Biosciences). Antibodies used were as follows: AKT (Cat#2920, Cell Signaling Technology), AKT-pS473 (Cat#4060, Cell Signaling Technology), ACTIN (Cat#MAB1501, Millipore), IRDye 800CW goat anti-rabbit IgG (Cat#926-32211, LI-COR), and IRDye 680RD goat anti-mouse IgG (Cat#926-68070).

For qPCR, total RNA was isolated using the RNeasy kit (Qiagen). RNA was reverse-transcribed to cDNA using iScript cDNA synthesis kit (BIO-RAD). Semiquantitative real-time PCR analysis was performed using fast SYBR green (Applied Biosystems). Relative expression levels were determined by normalizing each CT values to POLR2A using the ΔΔCT method. The sequence for the primers used in this study was as follows. IPPK-fw: 5'-AATGAATGGGGGTACCACGG-3', IPPK-rv: 5'-AACTTCAGAAACCGCAGCAC-3'; MINPP1-fw: 5'-AGCTACTTTGCAAGTGCCAG-3', MINPP1-rv: 5'-TGCATGACCAAACTGGAGGA-3'.

Knockout cells were generated using the LentiCRISPR system as described in Sanjana et al., 2014(*57*). gRNAs against IPPK and MINPP1 were expressed from LentiCRISPRv2 (kind gifts from Feng Zhang; Addgene plasmids no. 49535 and no. 52961) by transfection of HEK293T cells with 1µg DNA using jetPRIME. The following gRNA target sequences were used: IPPK gRNA 5'-TCGGCCGGTGCTCTGCAAAG-3', MINPP1 gRNA 5'-ATCCAGTCCGCGTACCACAA-3'. Following transfection, cells were selected with puromycin, propagated, and screened for loss of target protein by qPCR. DNA sequencing of PCR products confirmed insertions or deletions leading to interrupted sequencing reactions. Pools of knockout cells were used to avoid clonal variation. HEK293T cells transfected with empty vector were used as control.

Sample preparation for LC-MS analysis

10 µg of mTORC2 I- site mutants I2, I3 and A- site mutants A3, A4, and A5 were dissolved in 50 µl digestion buffer (1% sodium deoxycholate (SDC), 0.1 M TRIS, 10 mM TCEP, 15 mM chloroacetamide (CAA), pH = 8.5) using vortexing for trypsin digestion. For endoproteinase GluC and chymotrypsin digestion, the same protein aliquots were dissolved in 20 ul of a digestion buffer consisting of 1 M urea, 0.1 M ammoniumbicarbonate, 10 mM TCEP and 15 mM CAA. Samples were either incubated for 10 min at 95°C (trypsin) or 1h at 37°C (GluC and chymotrypsin) to reduce and alkylate disulfide bonds. Protein aliquots were digested over night at 37°C by incubation with sequencing-grade modified trypsin, GluC and chymotrypsin (all 1/50, w/w; Promega), respectively. Then, the peptides were cleaned up using iST cartridges (PreOmics, Munich) according to the manufacturer's instructions. Samples were dried under vacuum and dissolved in LC-buffer A (0.1 % formic acid) at a concentration of 0.05 µg/ul.

Targeted PRM-LC-MS analysis to confirm presence of mutations

To enhance the sensitivity of the LC-MS analysis, a label-free targeted LC-MS approach was carried out. Therefore, three lists of peptides considering the cleavage specificity of the three proteases used and containing all mutation sites was generated. The peptide sequences were imported into the Skyline (version 20.1 (https://brendanx-uw1.gs.washington.edu/labkey/project/home/software/Skyline/begin.view) to generate a mass isolation list of all doubly and triply charged precursor ions for each protease. These were then loaded into a Q-Exactive plus LC-MS platform and analyzed using the following settings; The setup of the μRPLC-MS system was as described previously(*58*). Chromatographic separation of peptides was carried out using an EASY nano-LC 1000 system (Thermo Fisher Scientific), equipped with a heated RP-HPLC column (75 μm x 30 cm) packed in-house with 1.9 μm C18 resin (Reprosil-AQ Pur, Dr. Maisch). Peptides were analyzed per LC-MS/MS run using a linear gradient ranging from 95% solvent A (0.15% formic acid, 2% acetonitrile) and 5% solvent B (98% acetonitrile, 2% water, 0.15% formic acid) to 45% solvent B over 60 minutes at a flow rate of 200 nl/min. Mass spectrometry analysis was performed on Q-Exactive plus mass spectrometer equipped with a nanoelectrospray ion source (both Thermo Fisher Scientific). Each MS cycle consisted of one MS1 scan followed by high-collision-dissociation (HCD) of the selected precursor ions in the isolation mass lists. Total cycle time was approximately 2 s. For MS1, 3e6 ions were accumulated in the Orbitrap cell over a maximum time of 50 ms and scanned at a resolution of 35,000 FWHM (at 200 m/z). MS2 scans were acquired at a target setting of 3e6 ions, accumulation time of 110 ms and a resolution of 35,000 FWHM (at 200 m/z). The normalized collision energy was set to 27%, the mass isolation window was set to 0.4 m/z and one microscan was acquired for each spectrum.

The acquired raw-files were converted to the mascot generic file (mgf) format using the msconvert tool (part of ProteoWizard, version 3.0.4624 (2013-6-3)). Using the MASCOT algorithm (Matrix Science, Version 2.4.1), the mgf files were searched against a decoy database containing normal and reverse sequences of the predicted SwissProt entries of Homo sapiens (www.ebi.ac.uk, release date 2019/12/09), the mTOR and Rictor mutations and commonly observed contaminants (in total 41,556 sequences for Homo sapiens) generated using the SequenceReverser tool from the MaxQuant software (Version 1.0.13.13). The precursor ion tolerance was set to 10 ppm and fragment ion tolerance was set to 0.02 Da. The search criteria were set as follows: full tryptic specificity was required (cleavage after lysine or arginine residues unless followed by proline), 3 missed cleavages were allowed, carbamidomethylation (C), was set as fixed modification and oxidation (M) as a variable modification. Next, the database search results were imported to the Scaffold Q+ software (version 4.3.2, Proteome Software Inc., Portland, OR) and the protein false identification rate was set to 1% based on the number of decoy hits. Specifically, peptide identifications were accepted if they could be established at greater than 97.0% probability to achieve an FDR less than 1.0% by the scaffold local FDR algorithm. Protein identifications were accepted if they could be established at greater than 65.0% probability to achieve an FDR less than 1.0% and contained at least 1 identified peptide. Protein probabilities were assigned by the Protein Prophet program(*59*)). Proteins that contained similar peptides and could not be differentiated based on MS/MS analysis alone were grouped to satisfy the principles of parsimony. Proteins sharing significant peptide evidence were grouped into clusters. Finally, a spectral library (*.blib) was generated from the assigned MS/MS spectra and imported to Skyline together with the acquired raw data files. Only precursor ions confidently identified by database searching and present in the spectral library were employed for quantitative analysis. Quantitative result reports were further analyzed by Microsoft Excel and PRISM (GraphPad Software, San Diego, US).

EM Sample preparation and data collection

Different conditions were screened for mTORC2 in the presence and absence of substrates (Extended Data Figure 2). For all conditions, freshly thawed mTORC2 aliquots were used to prepare samples with an mTORC2 concentration of 0.37 mg/mL. Shortly before grid preparation, the samples were diluted to reach a final mTORC2 concentration of 0.12 mg/mL.

For each grid, a small piece of continuous carbon was floated on top of the sample for one minute(*60*). The carbon was then picked with a Quantifoil R2/2 holey carbon copper grid (Quantifoil Micro Tools), which was swiftly mounted in a Vitrobot (Thermo Fischer Scientific) whose chamber was set to 4°C and 100% humidity. 5 µL of buffer was then added on top of the grid on the side showing the carbon covered with particles, which was immediately blotted with a setting of 0 to 6 seconds blotting time and rapidly plunge-frozen in a mixture 2:1 of propane:ethane (Carbagas)(*61*).

Data were collected using a Titan Krios (Thermo Fisher Scientific) transmission electron microscope equipped with either a K2 Summit direct electron detector (Gatan), a K3 direct electron detector (Gatan) or a Falcon 3EC direct electron detector (Thermo Fisher Scientific) using either EPU (Thermo Fisher Scientific) or SerialEM(*62*) (Extended Data Fig. 2). Cameras were used in counting and/or super-resolution mode. During data collection, the defocus was varied between -1 and -3 µm and four exposures were collected per holes. Stacks of frames were collected with a pixel size of 0.84 Å/pixel and a total dose of about 70 electrons/Å2.

Data processing

For all datasets, the initial processing was done in similar fashion. First, the stacks of frames were aligned and dose-weighted using Motioncor2(*63*). GCTF(*64*) was used to estimate the contrast transfer function (CTF) of the non-dose weighted micrographs. After a selection of good micrographs using both the quality of the power spectra and the quality of the micrographs themselves as criteria, particles were picked using batchboxer from the EMAN1.9 package(*65*) using particle averages from manually picked particles as references. Particles were extracted using Relion3.0(*66*), followed by two rounds of 2D classification using cryoSPARCv2 (Structura Biotechnology Inc.)(*67*) (Extended Data Table 2). The first reference was generated by ab-initio reconstruction using cryoSPARCv2. Good particles from 2D classification were then used for a homogeneous 3D refinement followed by non-uniform refinement using cryoSPARCv2. Two masks were then generated manually around each half of the pseudo-dimeric mTORC2 using UCSF Chimera(*68*) and two focused refinements around each half of the complex using cryoSPARCv2 were performed using those masks. For the dataset 1 which contained ΔPH-Akt1, the resolution was further improved by performing Bayesian particle polishing(*69*) followed by CTF refinement using Relion3.1. Those particles were again subjected to a round of non-uniform refinement and local refinement using cryoSPARC v2. For each reconstruction, the maps were sharpened using phenix.auto_sharpen(*70*) or were transformed to structure factors using phenix.map_to_structure_factors(*71*) and sharpened in COOT(*72*).

Further 3D classifications without alignment for local structural variability close to the catalytic center were performed using the particles from the datasets containing the purified Akt1 and, independently, the ones from the dataset with ΔPH-Akt1 using Relion3.0(*66*) and using a mask manually created in UCSF Chimera(*68*). After classification, the particles were used for refinement using cryoSPARCv2 (Structura Biotechnology Inc.). To compare the density of the sample with and without ATPγS, the final density (Volume A) was filtered to 4.2 Å and compared to the density without ATPγS (Volume F). Difference density was calculated using UCSF ChimeraX(*73*)

Modelling and docking

First, mTOR and mLST8 models were taken from the EM structure of mTORC2 (PDB : 5ZCS(*28*)) and each fold was rigid-body fitted into the better half of the density. Minor changes in mTOR conformation were done manually to fit the density, then Rictor and SIN1 were manually built de novo using COOT(*72*). Map quality enabled direct model building for structured regions, lower resolution density provided connectivity information for assigning and linking regions of Sin1 and Rictor as shown in Extended Data Fig. 5c and 6b. The second half of mTORC2 was made by copying and rigid-body fitting each chain of the first half in the second one. Finally, the structure of either one-sided or two-sided mTORC2 were refined using phenix.real_space_refine(*70*) (Extended Data Table 1), using Ramachandran and secondary structure restraints. As the horns of mTOR were flexible and their local resolution were significantly lower, additional reference restraints were applied, using PDB: 6BCX(*31*) as reference. The model was then validated by comparing the FSCs calculated for the experimental density and the models (Extended Data Fig. 3). In addition, both the half and full structure were also refined in their respective half map (half map 1) and the FSC of this structure against the same half map (half map 1), the other half (half map 2) and the full map were compared. The similarity of the curves shows that the structure was not overfitted(*74*).

Ligand identification via mass spectrometry

Inositol hexakisphosphate (InsP6) (Sigma-Aldrich) was directly dissolved in 10 mM ammonium acetate (pH8.5) and diluted to 50 µM. mTORC2 in cryo-EM buffer was buffer exchanged and concentrated in 10 mM ammonium acetate (pH8.5) using an Amicon Ultra 0.5 mL – MWCO 100kDa. The concentrated complex was mixed with an equal volume of Phenol at pH8, thoroughly vortexed for 30 seconds and incubated at room temperature for 30 minutes. The tube was then centrifuged for 5 min at 15000 xg. The aqueous phase was then used for mass spectrometry. A sample containing only buffer and no protein was subjected to the same treatment for reference. The samples were then mixed with four volumes of injection buffer (90% Acetonitrile, 9% Methanol, 50 mM ammonium acetate (pH7) and directly injected using a Hamilton syringe in a Synapt G2-SI HDMS (Waters) in negative mode and using the T-Wave IMS.

Figure generation

All density and structure representations were generated using UCSF ChimeraX(*73*). Difference densities were calculated in ChimeraX using the "volume subtract" command. Local resolutions were estimated using cryoSPARC v2 (Structura Biotechnology Inc.). The electrostatic surface representation of Rictor was generated using APBS (Adaptive Poisson–Boltzmann Solver(*75*)). Multiple sequence alignment was performed using Clustal Omega(*76*) and visualized with Espript(*77*). Conservation analysis was done with AL2CO(*78*) and visualized in UCSF ChimeraX(*73*).

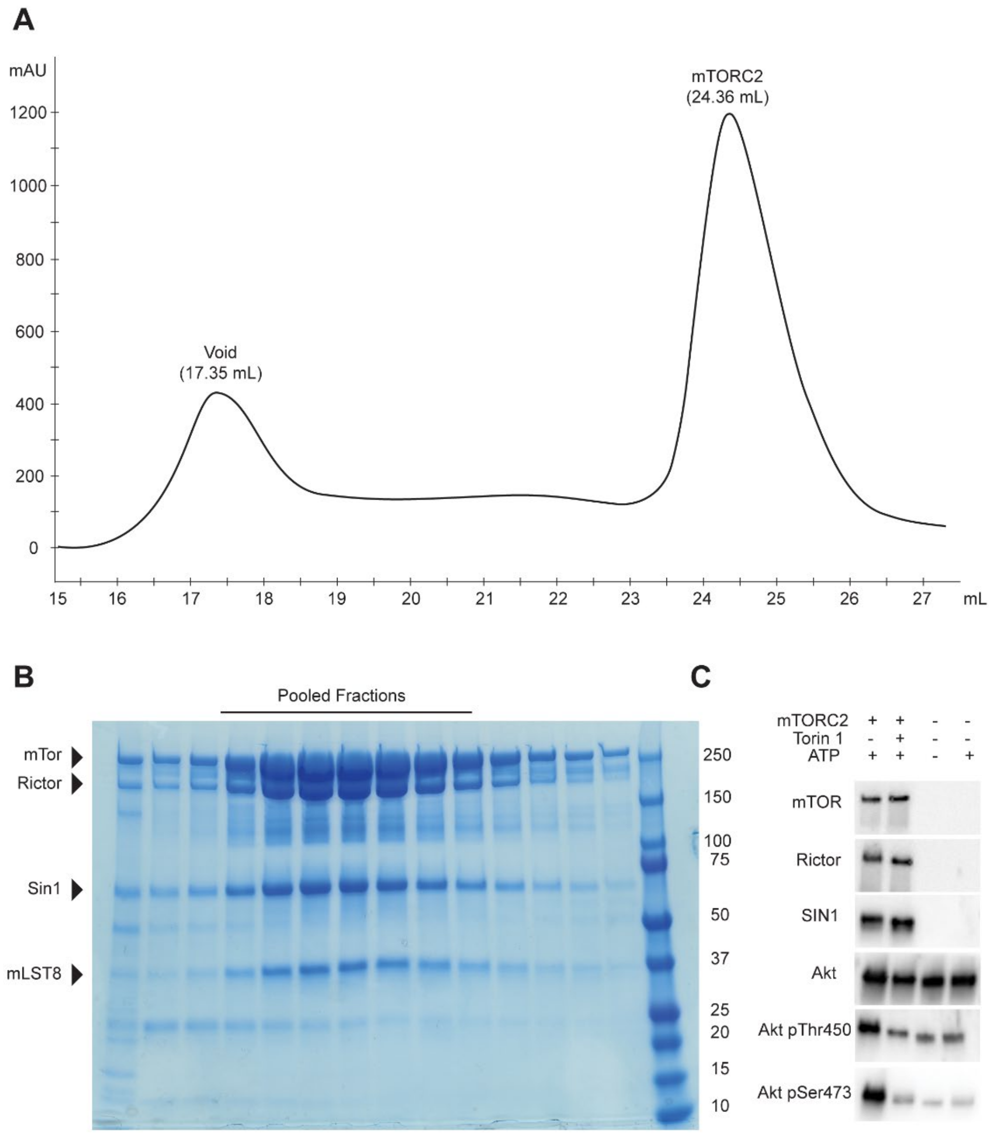


**Fig. S1.**

**mTORC2 purification and characterization.** (**A**) Size exclusion elution profile of mTORC2 from a custom made Superose 6 Increase 10/600 GL column at a flow rate of 0.1 mL/min. The void and the mTORC2 peaks are indicated. (**B**) SDS-polyacrylamide gel of the fractions of the size exclusion chromatography. Fractions pooled for further analysis are indicated. (**C**) Kinase activity assay of purified mTORC2 using Akt1 as a substrate. Western blots showing the phosphorylation state of Akt1 in the presence and absence of mTOR inhibitor Torin1, ATP and mTORC2. Akt1 phosphorylation is detected by phospho-specific antibodies as described in Methods. Unprocessed blots are shown in the Supplementary Data.

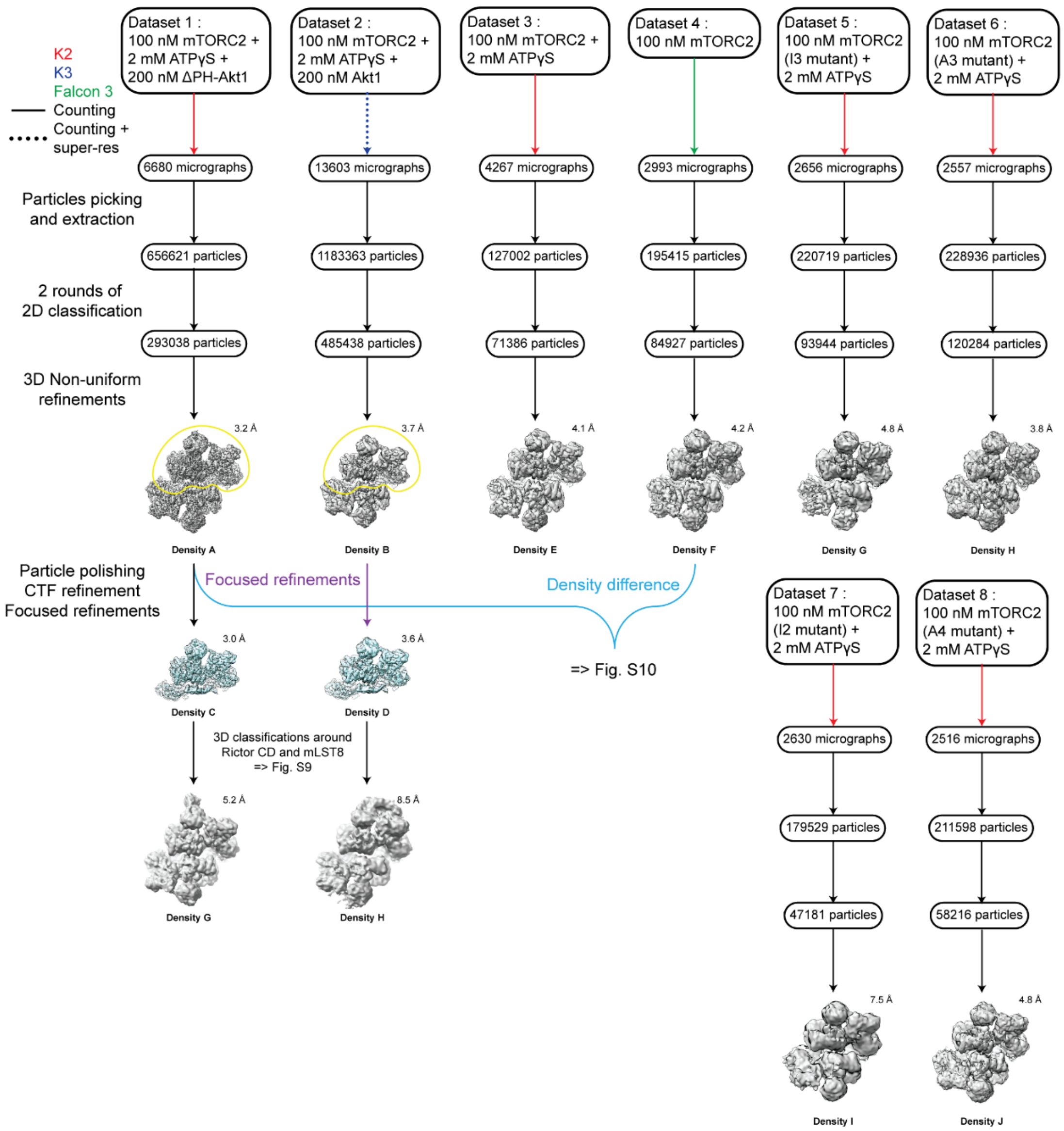


**Fig. S2.**

**Cryo-EM data processing scheme.** Four datasets were collected with different conditions: with full-length Akt1 (dataset 2), ΔPH-Akt1 (dataset 1) and without Akt1 (dataset 3) in the presence of ATPγS, and dataset 4 without addition of ATPγS. Dataset 1 reached the highest resolution and was used for further processing using CTF parameter refinement and Bayesian particle polishing to reach the highest resolution of this study, Density C, with a resolution of 3.0Å. Both datasets 1 and

2 were used for 3D classifications in the proximity of mLST8 after a refinement focused on the better half. Four further datasets (datasets 5-8) were collected for mTORC2 containing mutant variants of mTOR and Rictor, respectively, to analyze the impact of mutations.

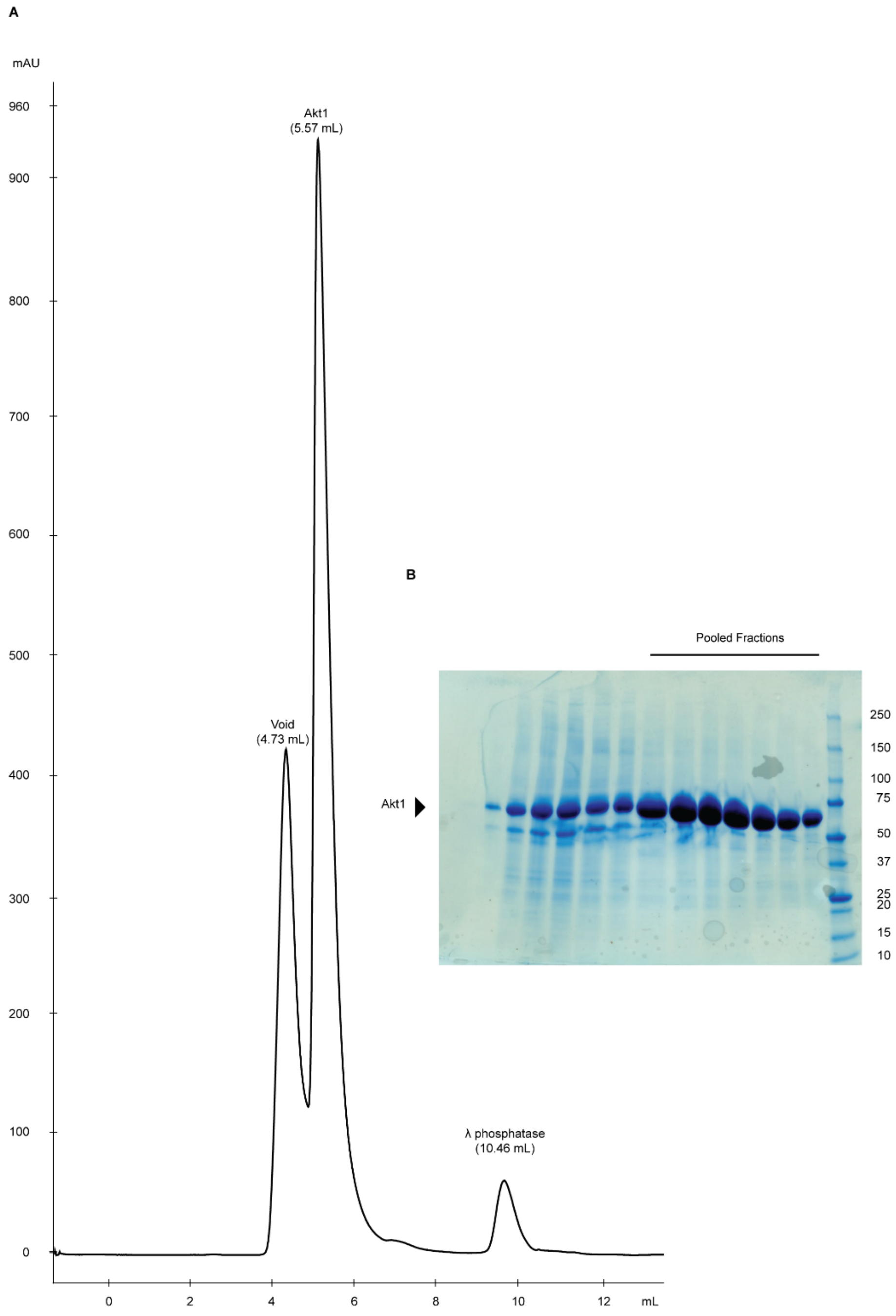


**Fig. S3.**

**Purification of Akt1.** (**A**) Elution profile of Akt1 after dephosphorylation by λ-phosphatase in size exclusion chromatography on Superdex 75 Increase. The void, the Akt1 and λ-phosphatase peaks are indicated. (**B**) SDS-polyacrylamide gel of the fractions of the size exclusion elution. Fractions pooled for the final sample are indicated.

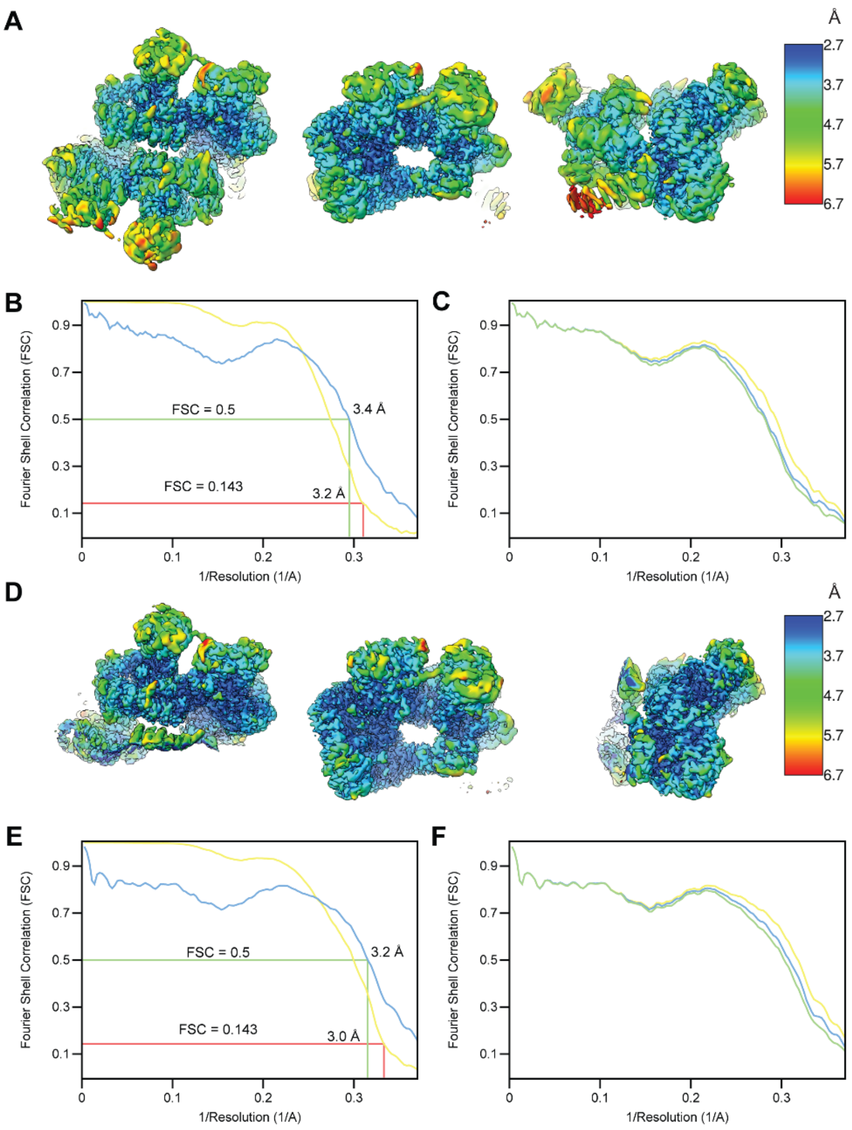


**Fig. S4.**

**Resolution of cryo EM reconstructions of mTORC2.** (**A**) Three views of local resolution heatmaps for non-uniform refinement (Density A). Local resolution varies between ~2.7 Å in the core of one half and 6-7 Å for the flexible region of the second half. (**B**) FSC curves calculated between the two half maps (yellow) and between the model and the non-uniform refinement map (blue). The overall resolution of the map is 3.2 Å (FSC=0.143) and is close to the 3.4 Å resolution of model versus map (FSC=0.5). The large local resolution differences (Panel A) are most likely responsible for the deviation between half-map and model-based resolution. (**C**) FSC curves calculated between the model refined in the half map 1 versus the half map 1 (blue), the half map 2 (green), and the full map (yellow). (**D**) Same as panel **A** but for the good half (Density C), which was used for model building and refinement. (**E**) and (**F**) similar to **B** and **C** respectively, but for maps calculated with a mask around the good half.

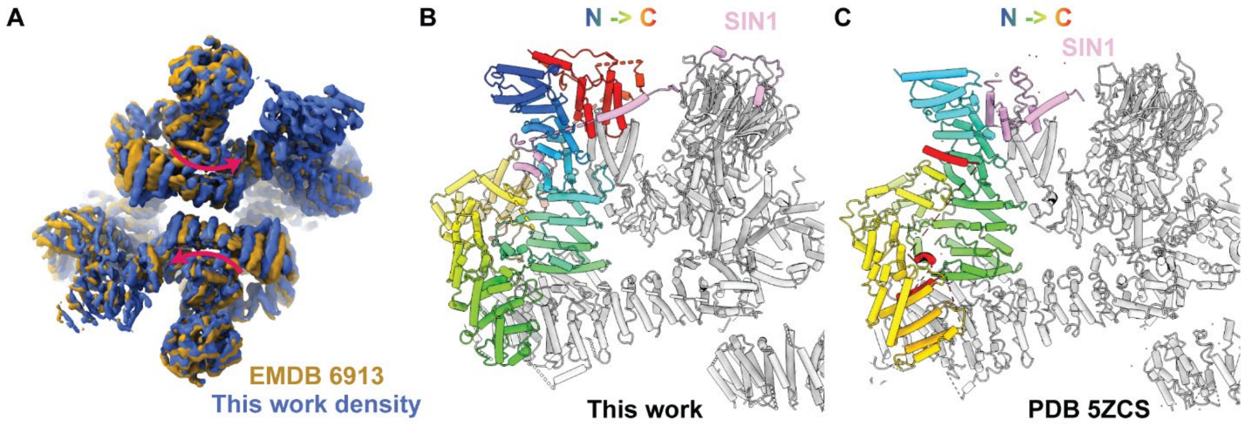


**Fig. S5.**

**Comparison of current and previous mTORC2 structures.** (**A**) Overlay of mTORC2 density A (blue) filtered down to 4.9 Å to match the resolution of an earlier mTORC2 reconstruction (EMDB 6913 *(28)*) showing the inward rotation of the FAT region in the current reconstruction. A similar mode of rotation is apparent when comparing to an earlier intermediate resolution structure of human mTORC2 (EMDB 3927 *(30)*). (**B**) and (**C**), Overview of Rictor and SIN1 topology in the current structure (B) and a previous mTORC2 model (PDB: 5ZCS *(28)*) (C). SIN1 is coloured in pink and Rictor is coloured by sequence from blue (N-terminus) to red (C-terminus).

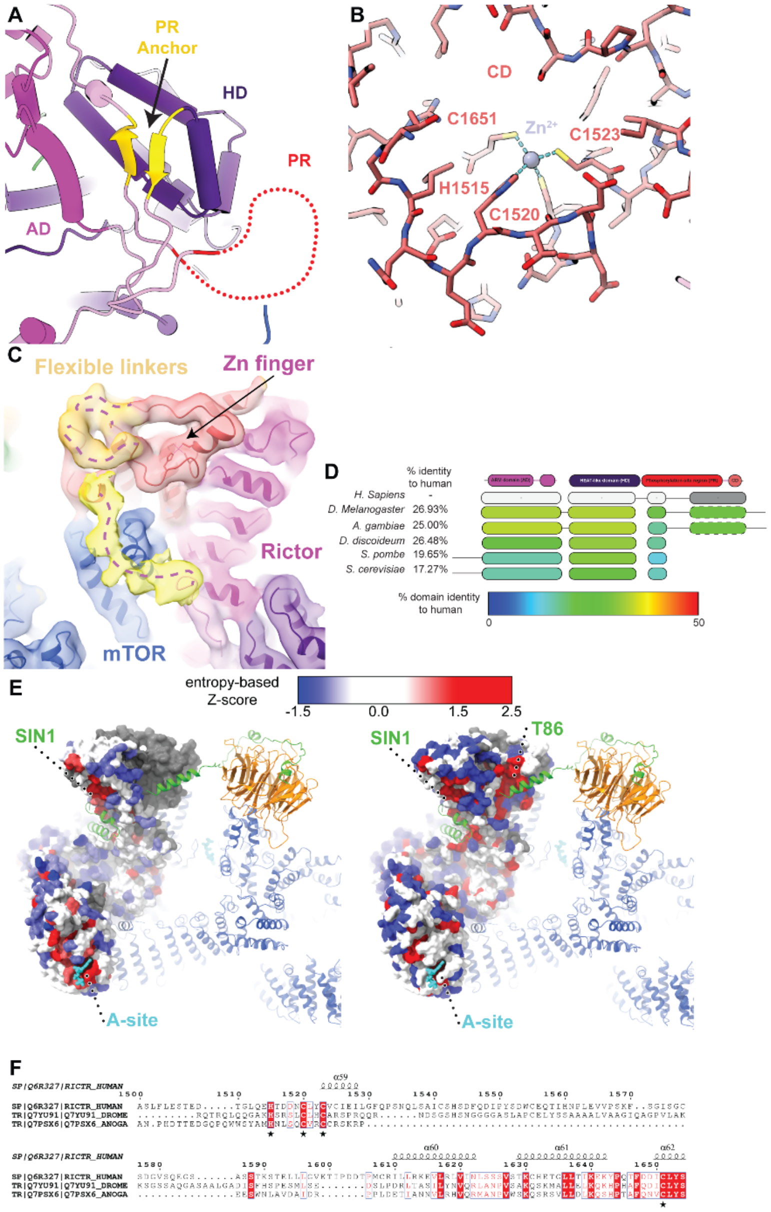


**Fig. S6.**

**PR anchor and $Zn^{2+}$ binding site in Rictor.** (**A**) Close-up view of the antiparallel-strands (gold) which are located in sequence before and after the PR of Rictor, anchoring the PR to the top of HD. (**B**) Close-up view of the Rictor CD zinc finger, which is part of the four-helix bundle and is formed by His1515, Cys1520, Cys1523, and Cys1651. (**C**) An overall mTORC2 reconstruction filtered to 6Å resolution reveals flexible linker segments (dashed line) and the connectivity in the

CD region. The map is shown at 4σ contour level and flexible linkers are highlighted in orange (3.5σ contour level). Another linker visible at lower contour level (shown at 2.5σ contour level in yellow) shows the connectivity to the preceding structured segment of Rictor. (**D**) Sequence conservation in Rictor. A multiple sequence alignment of Rictor orthologs is analysed by AL2CO *(78)* (entropy based conservation measure, independent counts) which shows four conserved blocks colored according to their respective percent identity relative to *H. sapiens* Rictor. The fourth block, corresponding to the end of the PR and the CD, is only conserved in metazoans. (**E**) Surface representation of Rictor coloured by residue conservation within eukaryotes (left) and metazoans (right). Blue represents poorly conserved residue, while red shows high conservation. Conserved patches are found around the A-site and the region interacting with SIN1. Grey regions are not found in all species compared. (**F**) Multiple sequence alignment of metazoan homologs of Rictor around the Zinc finger. The residues coordinating the zinc are underlined with a star.

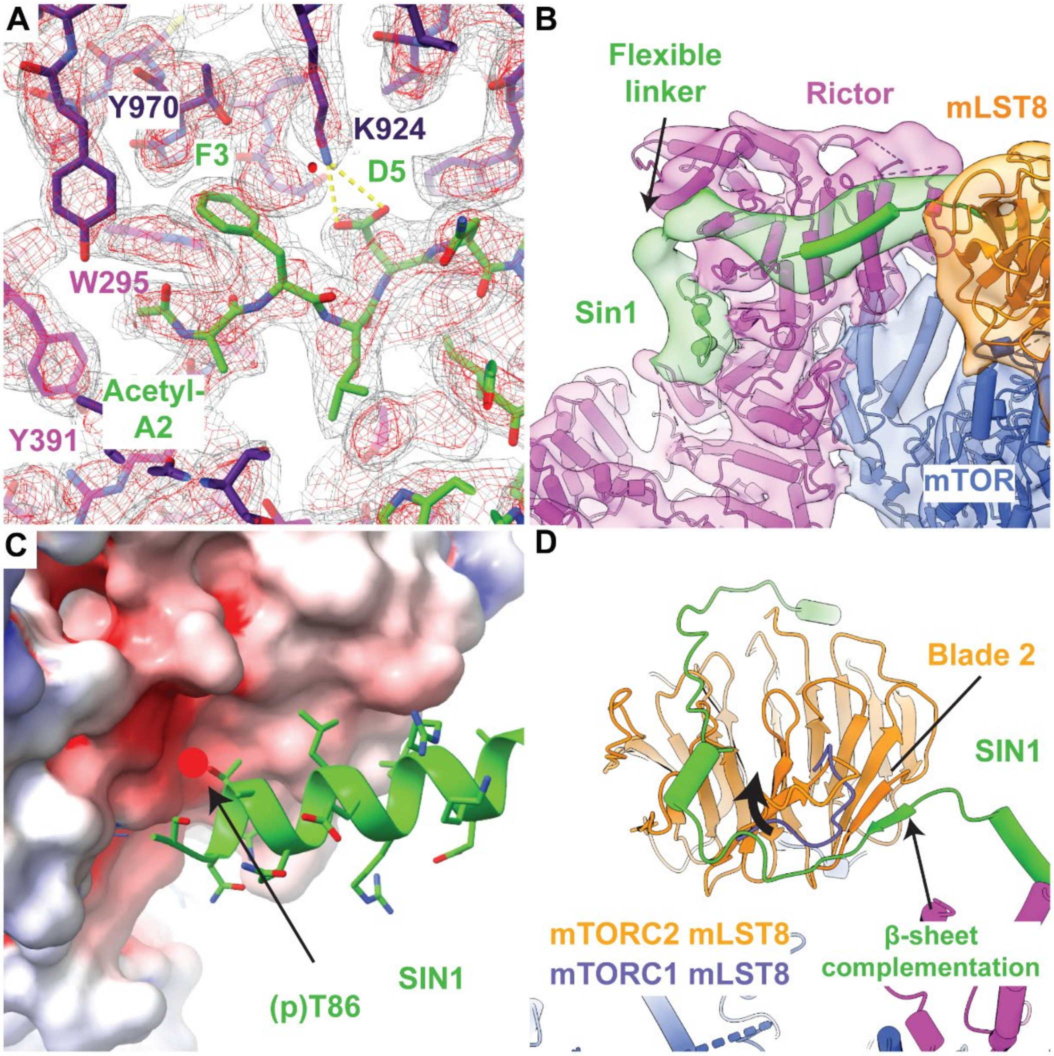


**Fig. S7.**

**Local structural features and interactions of SIN1.** (**A**) Close-up view of the N-terminal end of SIN1 (green), which is found deeply inserted between the AD (magenta) and the HD (dark magenta) of Rictor. Possible hydrogen bonds are indicated as a dashed yellow line. (**B**) Overview of the poorly ordered SIN1 linker between its N-terminal region and the traverse shown using a map filtered to 10Å resolution. (**C**) Close-up view of Thr86 of SIN1. Rictor is shown as a surface and coloured according to the electrostatic potential. Thr86 is inserted into a negatively charged pocket and phosphorylation (red circle) is likely incompatible with insertion into this pocket.

(**D**) Different conformations of the linker between sheet three and four of the third blade of mLST8 are observed in mTORC2 (orange) and mTORC1 (blue). SIN1 extends towards the second blade of mLST8 and interacts via β-strand complementation.

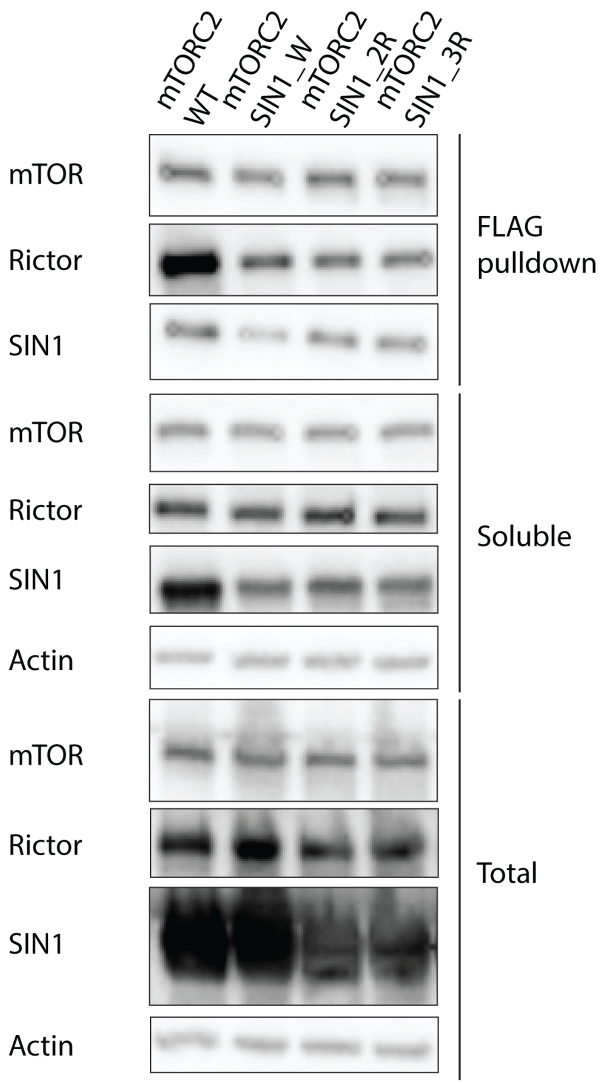


**Fig. S8.**

**Analysis of expression and mTORC2 integration of SIN1 variants.** Western blot from lysate and mTOR-based FLAG-bead pulldown of in insect cells recombinantly overexpressed mTORC2 WT and mTORC2 carrying variants of SIN1 with insertion of a tryptophan (mTORC2 SIN1_W), two consecutive arginines (mTORC2 SIN1_2R) and three consecutive arginines (mTORC2 SIN1_3R) at its processed N-terminus. Total lysate, soluble input protein after centrifugation and Flag-bead pulldown were blotted with immunodetection for mTOR, Rictor and SIN1. Actin was used as loading control. mTOR, Rictor and SIN1 expression levels are comparable in the WT and SIN1 N-terminal variants, but Rictor and SIN1 pull-down is lower in mutant variants than in mTORC2 WT.

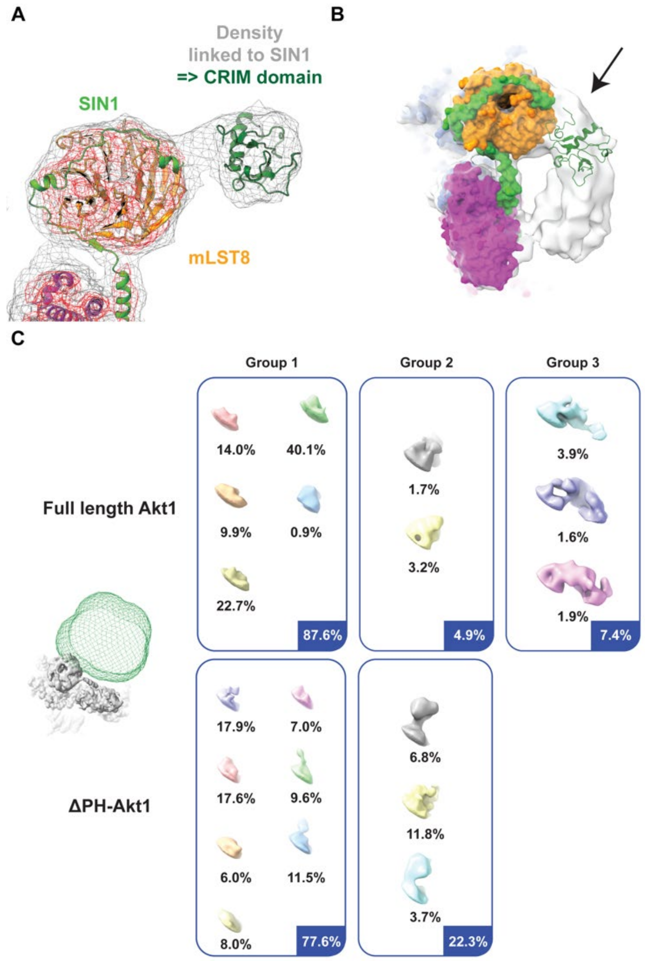


**Fig. S9.**

**Local structural features and substrate recognition by SIN1.** (**A**) Overview of SIN1 and mLST8 in density G (Fig. S2), shown at two different contour levels (red and grey), highlights the extra density found in the continuation of SIN1. (**B**) View of the structure above the catalytic site shown within density H (Fig. S2) showing the visible arch of extra density found between mLST8 and Rictor. The structure of the CRIM domain (PDB 5RVK) was fitted into the density, explaining part of the arch. The second part cannot be explained unambiguously. (**C**) Local classification without alignment around the SIN1 CRIM domain in density G (Fig. S2). Top view of 3D classes calculated with a mask around Rictor CD and mLST8. (show on the left in green relative to the high-resolution reconstruction). The top ten classes are calculated from datasets collected for samples containing full length Akt1, while the bottom ten classes are of samples containing ΔPH-Akt1. Classes showing small extra density, probably corresponding to the CRIM domain were put in group 2, while the classes showing a more “arch”-like structure were put in the group 3.

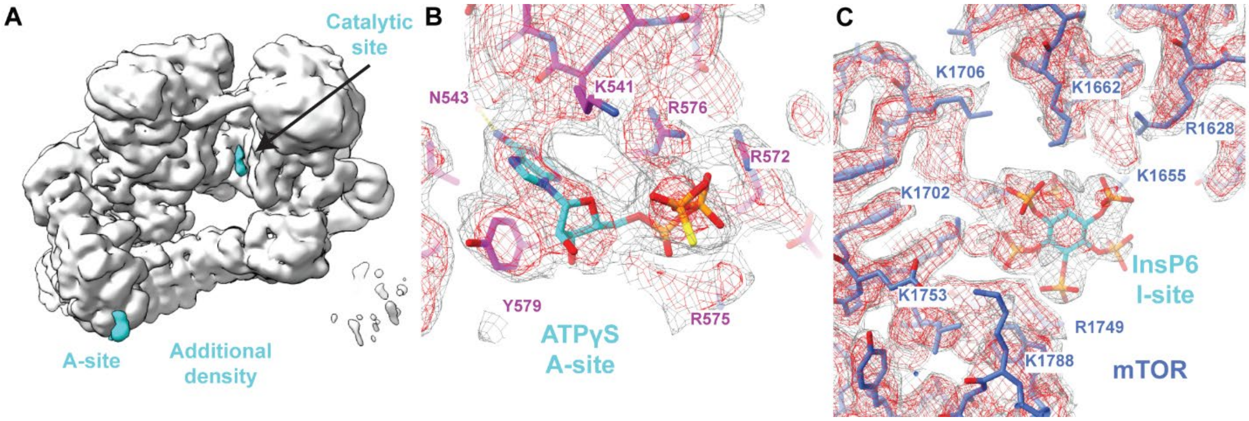


**Fig. S10.**

**Overview of mTORC2 ligand binding sites.** (**A**) Difference densities between the reconstructions in presence (Density A, low pass filtered to 3.9Å) and absence of ATPγS (Density D, showed in grey) are shown in light blue (shown at 4σ) in the A-site and the catalytic site, highlighting the similarity of differences in both site (**B**) Close-up view of ATPγS in the A-site in Rictor. (**C**) Close-up view of InsP6 bound to the I-site in the FAT domain of mTOR (blue). For panel (B), and (C), density for the focused refinement around one protomer is shown in mesh style at two contour levels (grey and red). (Density D).

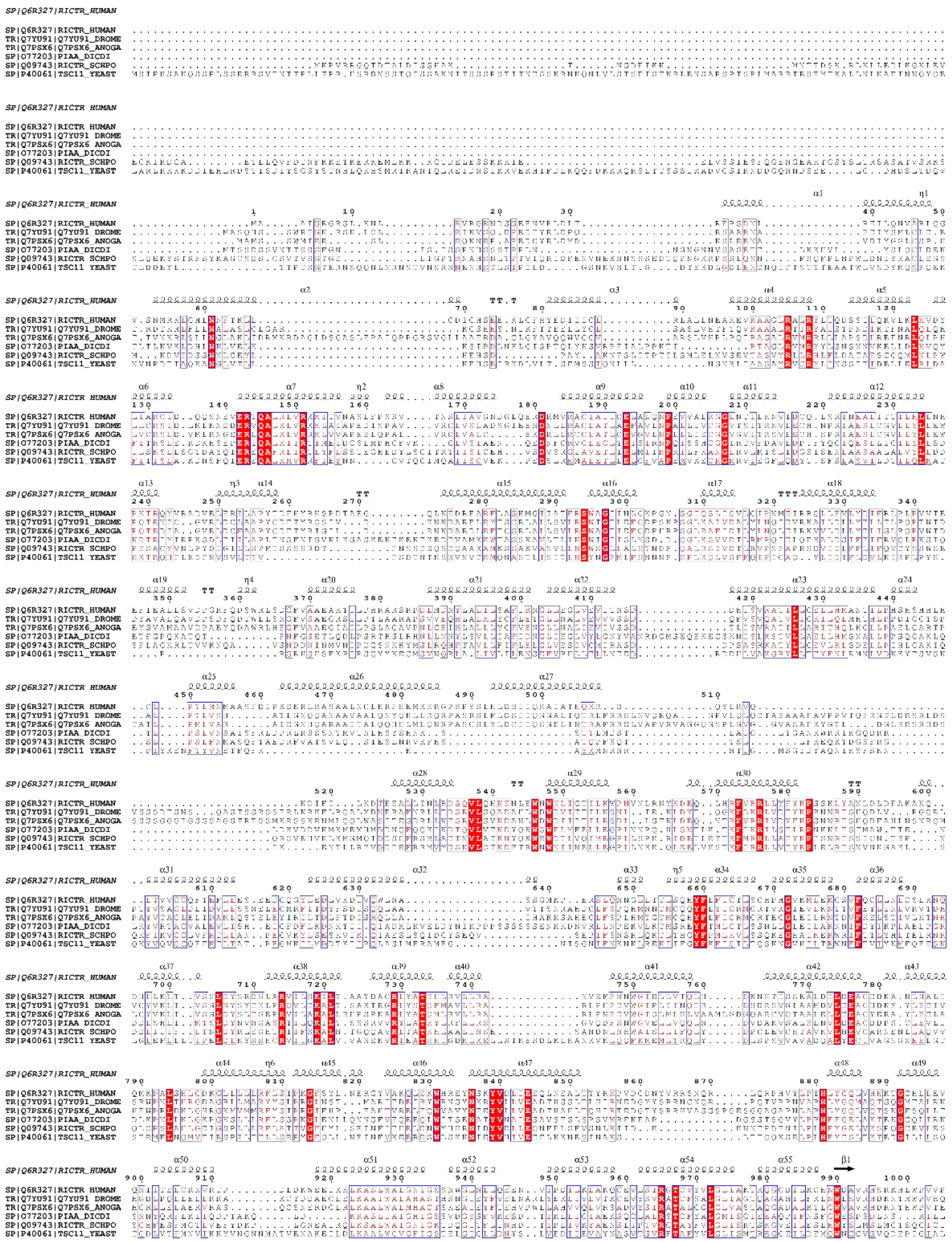

**Fig. S11.**

**Sequence conservation of Rictor.** Multiple sequence alignment of Rictor orthologs showing conservation and secondary structure. Blue boxes represent region of BLOSUM62 similarity score above 0.1, while the one highlighted in red are strictly conserved among the shown sequences.

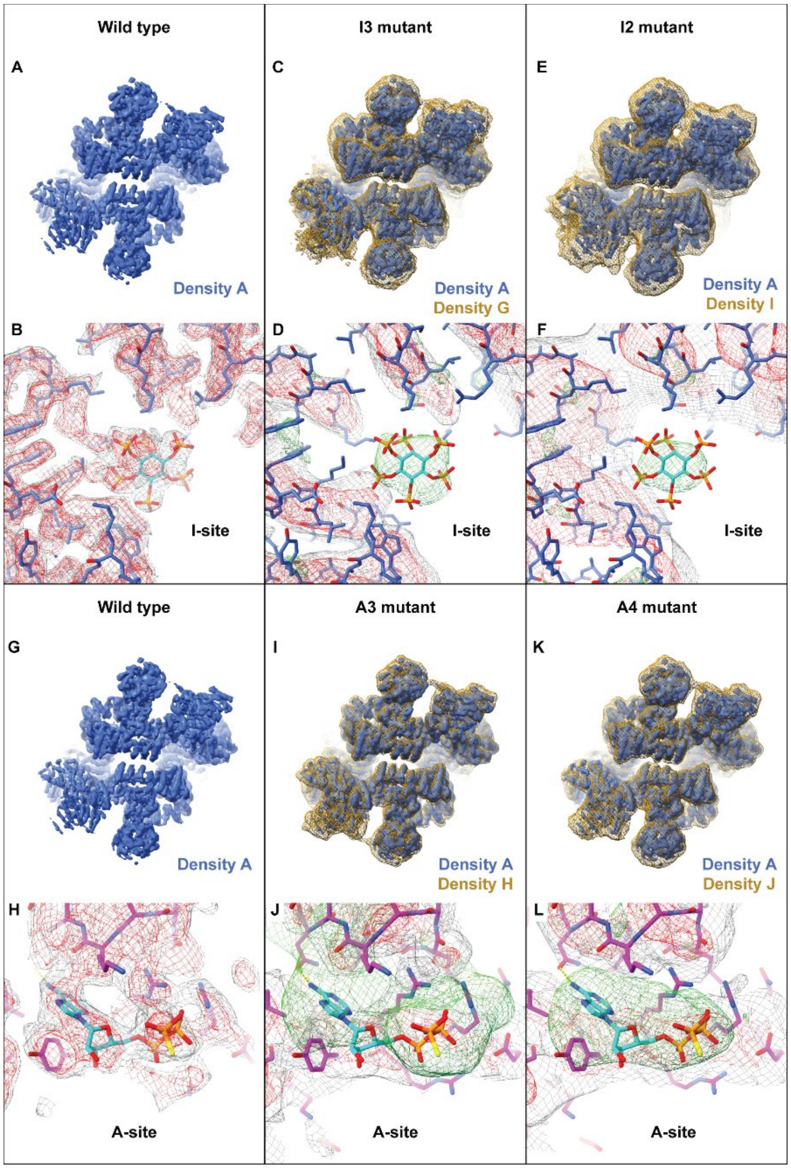


#### Fig. S12.

**Comparison of reconstructions of wild type mTORC2 and I-site or A-site variants.** (**A**) and (**G**): Overall cryoEM reconstruction of wt-mTORC2 in presence of ATPγS and ΔPH-Akt1 for the indicated variants of mTORC2. (**B**). Close-up view of the I-site in the wt-mTORC2. (**C**), (**E**), (**I**), and (**K**) Overlay of mTORC2 density A (blue) filtered down to 4.9 Å with mutant mTORC2 reconstruction. (**D**),(**F**). Close up-view of the I-site of the mTOR mutants (grey and red) with the difference density with the wt-mTORC2 (green). (**H**) Close-up view of the A-site in the wtmTORC2. (**J**),(**L**). Close up-view of the A-site of the mutant Rictor mutant (grey and red) with the difference density with the wt-mTORC2 (green).

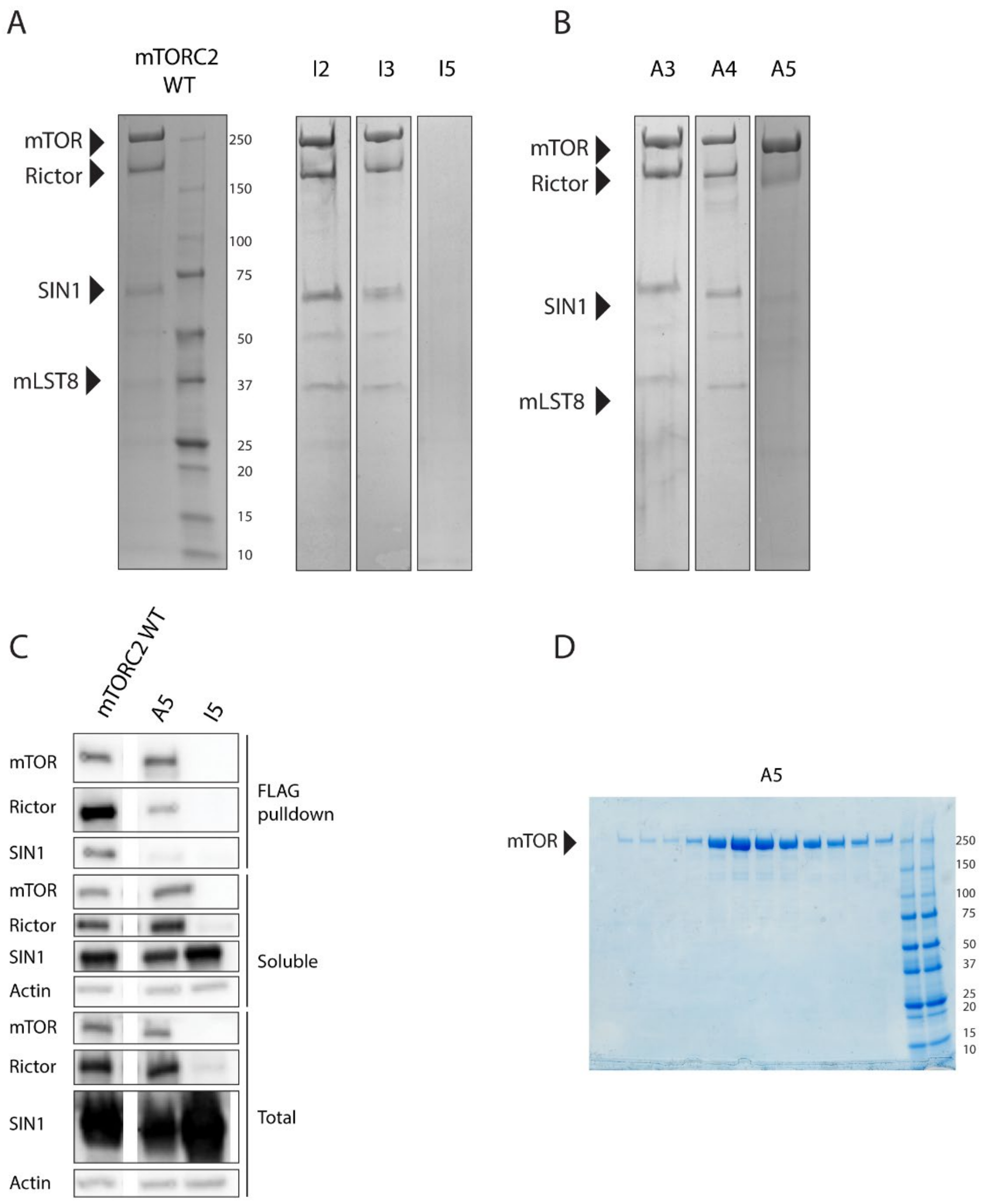


**Fig. S13.**

**Expression and assembly of A- and I- site mutants into mTORC2.** (**A**) Coomassie-stained SDS-polyacrylamide gel analysis of mTOR-based FLAG bead pulldown from recombinant overexpression in insect cells for mTORC2 WT and three variants with mutations in the I-site. mTOR mutants I2 and I3 assemble into mTORC2 complexes, for mutant I5, neither mTOR nor other mTORC2 subunits are pulled down. (**B**) Coomassie-stained SDS-polyacrylamide gel of a small-scale mTOR-based FLAG bead pulldown of three Rictor A-site mutants.

Mutants A3 and A4 assemble into mTOR complexes. Mutant A5 shows lower level of pulled down Rictor and SIN1. (**C**) Analysis of protein levels from recombinant overexpression in insect cells of mTORC2 components for mutant variants I5 and A5, which show no or partial complex assembly. Rictor mutant A5 shows comparable expression levels to WT in the total and input fractions but less SIN1 and Rictor are pulled down, indicative of defective complex assembly. For mTOR mutant I5, mTOR is not detected, Rictor levels are lower than WT, but SIN1 is present at WT levels. (**D**) SDS-polyacrylamide gel of size exclusion chromatography after up-scaled recombinant insect cell overexpression of mTORC2 with Rictor variant A5. Purification yielded a diminished Rictor content, indicating defective complex assembly.

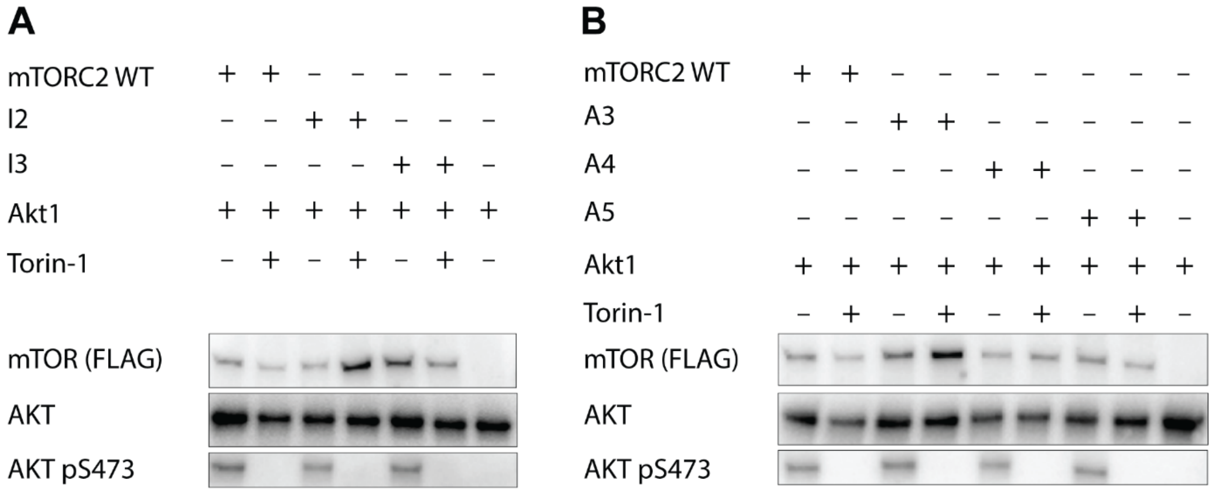


**Fig. S14.**

**Effect of A- and I- site mutations on mTORC2 activity in vitro.** In vitro kinase activity assay of purified mTORC2 for two I-site and three A-site variants (A5 purifies as only mTOR) compared to mTORC2 WT, using Akt1 as substrate and immunodetection of AKT pSer473 as readout for activity. Addition of Torin-1 is used for negative control. (**A**) Analysis of I-site variants I2 and I3, all variants display activity. (**B**) Analysis of A-site variants A3, A4 and A5 (purified with diminished Rictor content), all variants display activity.

A

| | Apparent Melting Point °C | | |
| --- | --- | --- | --- |
| Sample ID | Mean | SD | Replicates |
| mTORC2 WT | 44.6 | 0.428 | 5 |
| I2 | 42.8 | 0.354 | 5 |
| I3 | 44.4 | 0.339 | 5 |
| A3 | 44 | 0.563 | 5 |
| A4 | 44 | 0.443 | 4 |
| A5 | Not measurable | | |

B

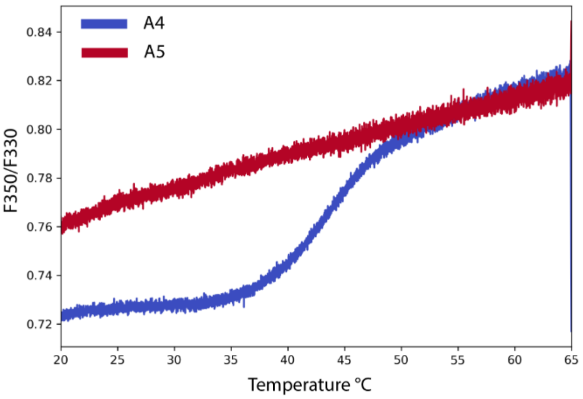


**Fig. S15.**

**Thermal stability of A- and I- site variants of mTORC2.** (**A**) Mean and SD of the mean of the melting points of mTORC2 WT and two I-site and two A-site variants, as recorded by nanoDSF measurements. (**B**) Plot of the ratio of fluorescence at wavelengths 350/330 nm plotted against increasing temperature to determine the Tm of the analyzed protein. Variant A5, which purifies with diminished Rictor content shows no defined melting point, in contrast to variant A4 that contains only one mutation less, indicating exposure of tryptophan residues already at low temperatures.

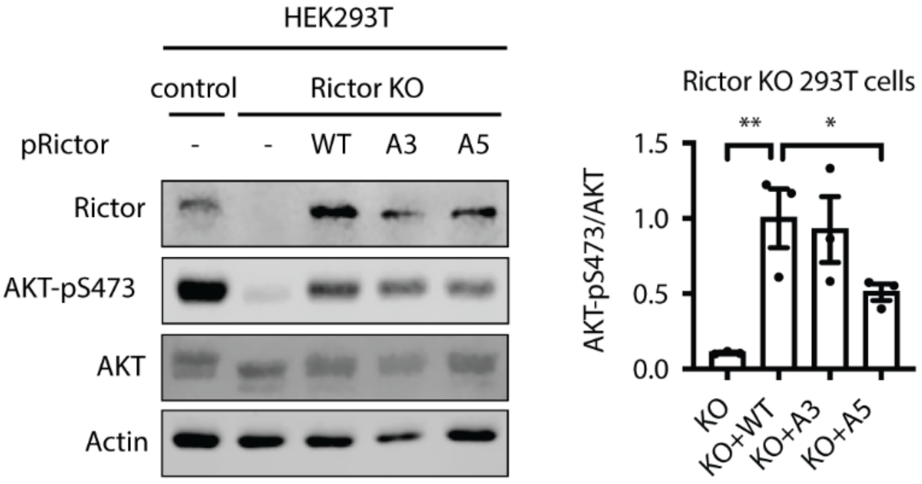


**Fig. S16.**

**In cell mTORC2 activity for mTORC2 A-site variants.** Immunoblots of lysates from control or Rictor knockout HEK293T cells complemented with Rictor-WT, Rictor variant A3, or Rictor variant A5. Cells were starved for serum for overnight and stimulated with 10% FCS and 100 nM insulin for 15 min. Actin serves as a loading control. For quantification, AKT-pS473 signals are normalized to total AKT signals. One-way ANOVA, $^{**}p<0.01$, $^{*}p<0.05$. N=3.

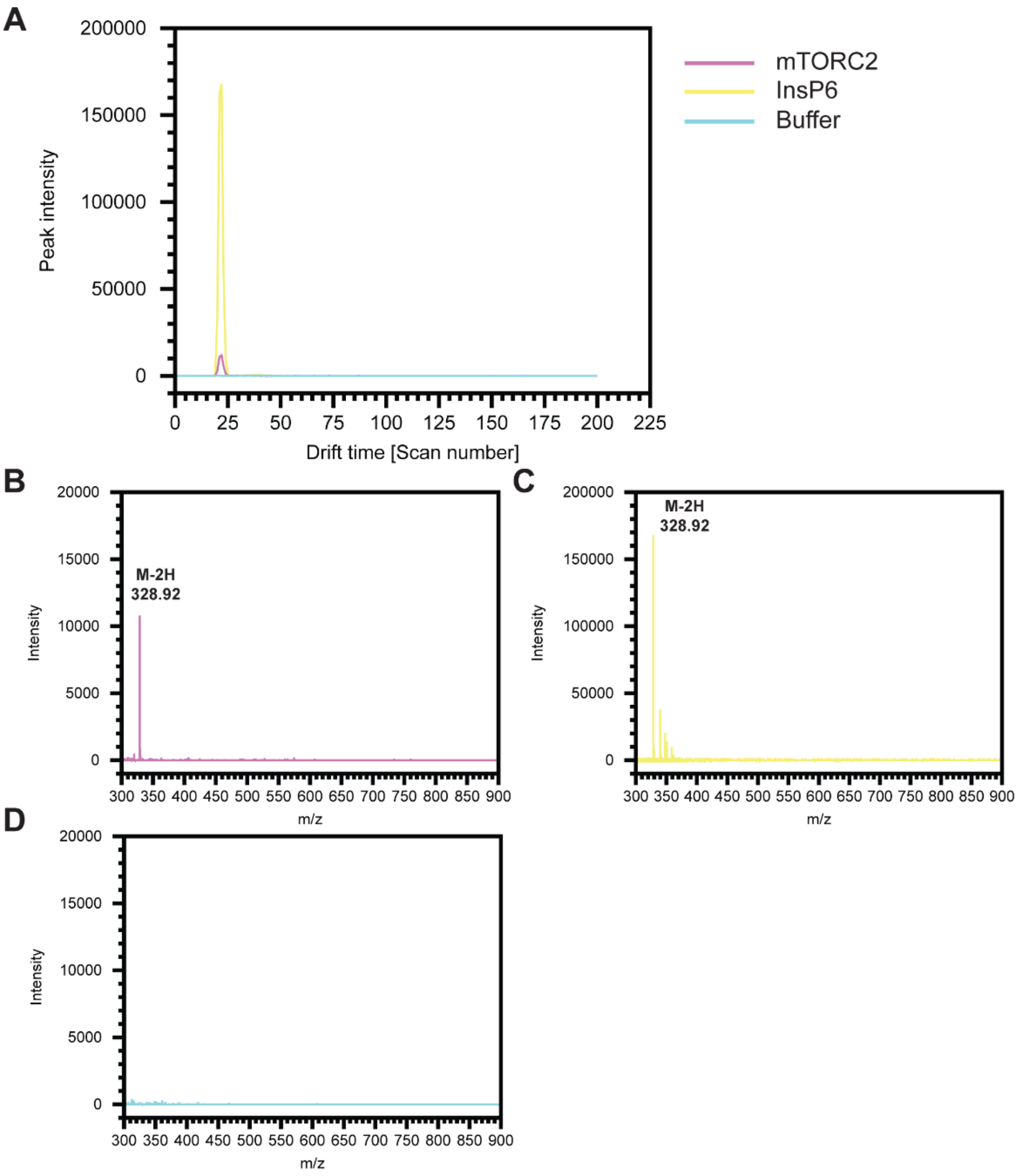


**Fig. S17.**

**Mass spectrometry analysis for ligand identification.** (**A**) Ion mobility chromatogram for the m/z 328.92 +/- 0.01 Da. The buffer control curve (blue) which was treated as the sample one (magenta) does not show any peak in this range, while the InsP6 reference (yellow) shows a strong peak around scan 22. The same peak was found in our sample (magenta) at the same drift time. (**B**) Mass spectra corresponding to the scan 22 in the ion mobility is shown between 300 and 900

Da. Peaks corresponding to the doubly charged InsP6 are observable in the mTORC2 sample (magenta) (**B**) and in the InsP6 reference sample (yellow) (**C**). No significant peaks are detected in the buffer control.

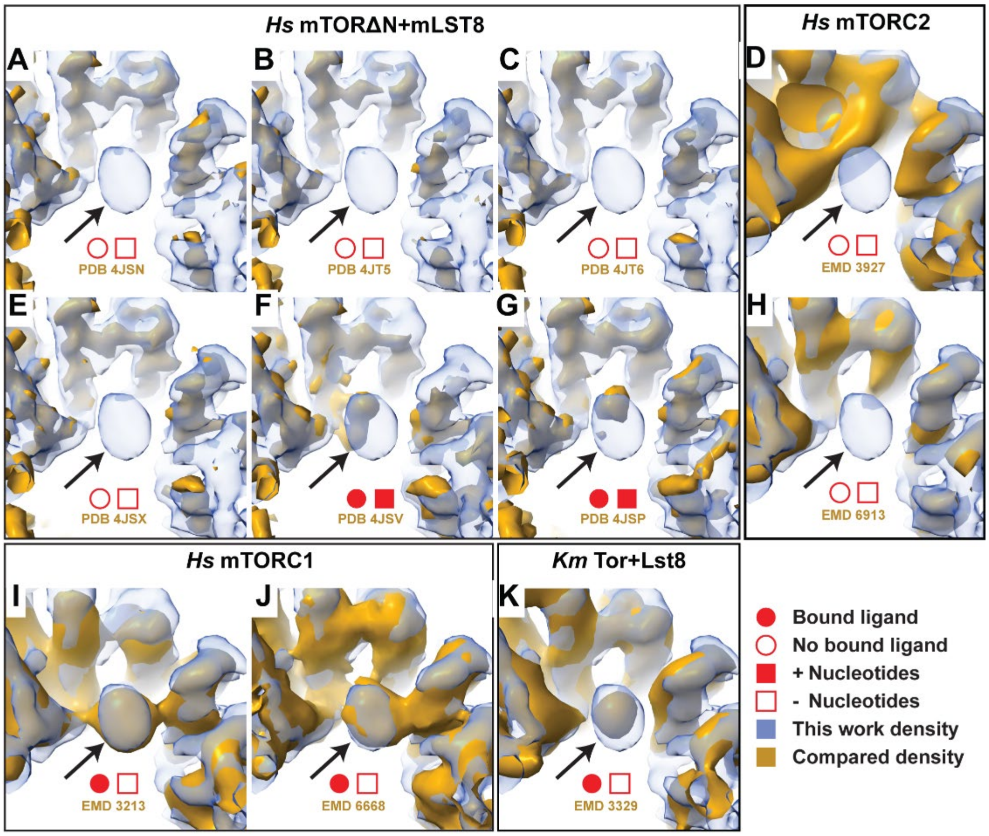


**Fig. S18.**

**Comparison of the I-site in published structures of mTOR complexes.** Close up view of the I-site (arrow) in the mTORC2 density (blue mesh) compared with published density maps (dark yellow). (**A-C**) and (**E-G**). Densities for crystal structures of human mTOR-mLST8. Extra density can be seen in the I-site in densities corresponding to PDB 4JSV and 4JSP panel F and G. These structures were obtained in presence of ATP analogs. (**A,B,C** and **E**) structures were obtained in absence of ATP or ATP analogs. (**D** and **H**), Comparison to human mTORC2 from EMD3927 and EMD 6913 in which no ligand binding is observed. (**I-K**) Densities from low and medium resolution cryoEM reconstructions which also show extra density in the I-site. The densities come from (**I**) human mTORC1 (EMD 3213) and from (**J**) human mTORC1(EMD 6668) and from (**K**) fungal Tor-Lst8 (EMD 3329). (**I-K**) Densities were obtained in absence of ATP analogs. Together these results suggest that InsP6 can be copurified presumably depending on cellular InsP6 concentration and specific purification conditions. Appearance of density in the I-site upon nucleotide analogue supplementation suggests the possibility of alternate binding of nucleotides to the mTOR I-site, when it is not occupied by Ins6P.

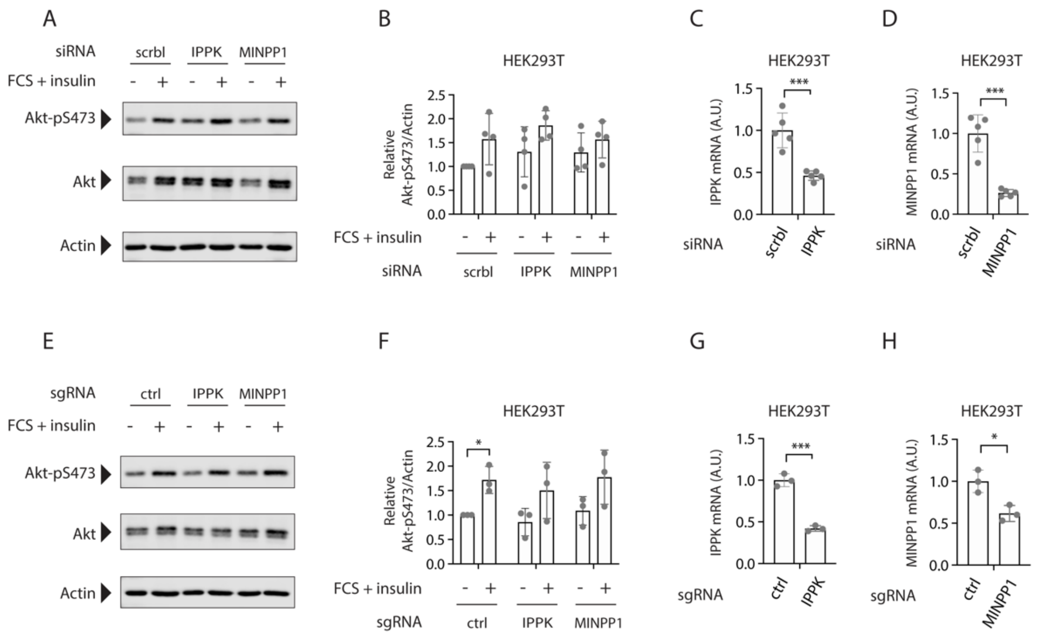


**Fig. S19.**

**Knock-down or knock-out of IPPK or MINPP1 does not alter mTOR-Akt signalling** (**A**) Western Blot showing Akt phosphorylation under starving and stimulating conditions. IPPK and MINPP1 knockdown cells were generated using HEK293T cells. n=3 (**B**) Quantification of western blot in A) and all biological replicates (n=3). (**C,D**) Knockdown validation via qRT-PCR analysis of IPPK and MINPP1 knockdown cells (n=5). (**E**) Western Blot showing Akt phosphorylation under starving and stimulating conditions. IPPK and MINPP1 knockout cells were generated using HEK293T cells. n=2. (**F**) Quantification of western blot in D) and all biological replicates (n=2). (**G,H**) Knockout validation via qRT-PCR analysis of IPPK and MINPP1 knockouts in HEK293T cells (n=2). Sequencing results of PCR fragments including the sgRNA binding site were interrupted, which suggests successful Cas9 activity and knockout of IPPK and MINPP1.

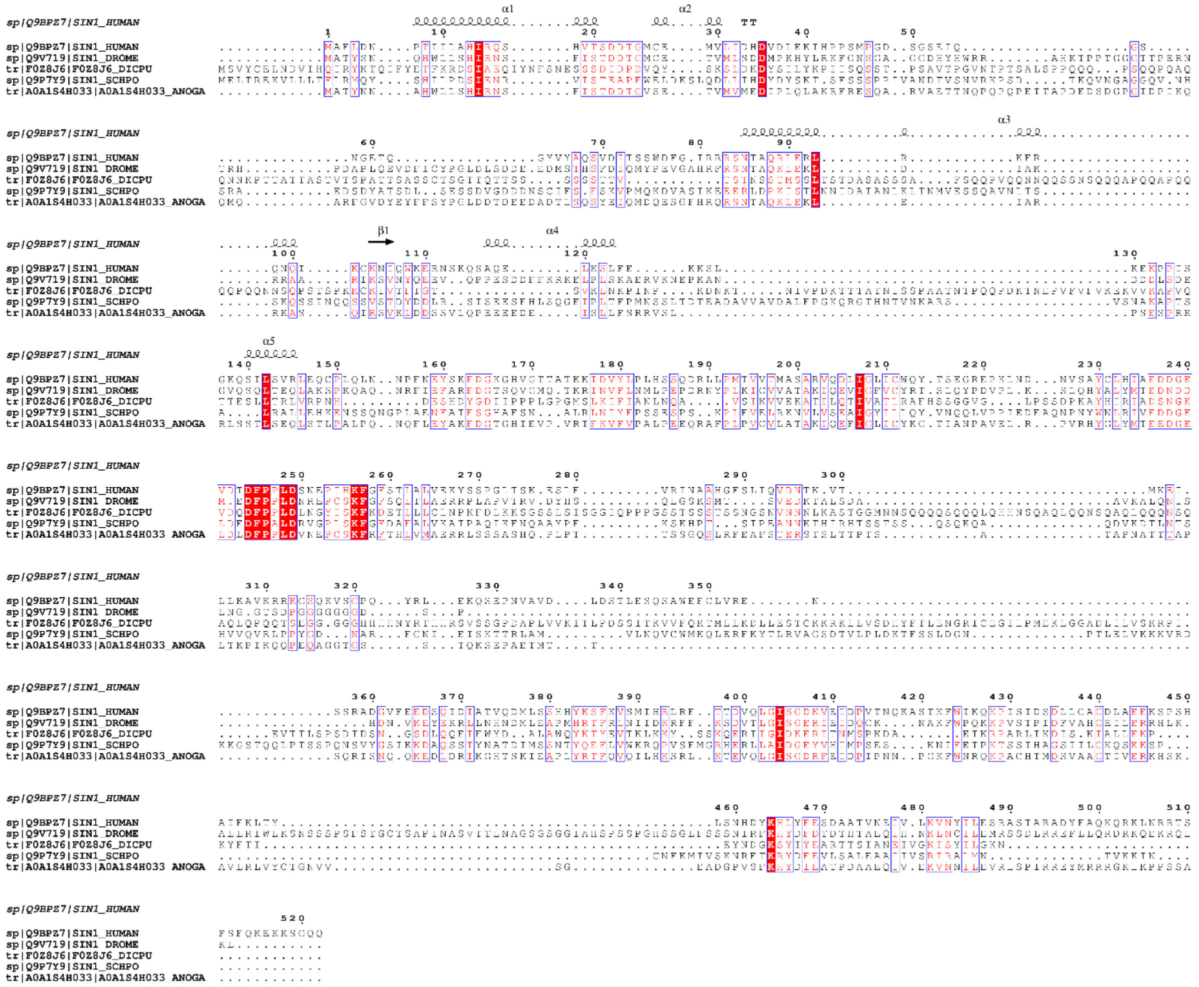

**Fig. S20**

**Sequence conservation of SIN1.** Multiple sequence alignments of SIN1 homologs showing conservation and secondary structure. The sequence of the S. cerevisiae SIN1-homolog Avo1 was omitted for clarity as it contains several large insertions.

**Table S1.**

| ID | Mutant type | Mutations | Primers |
| --- | --- | --- | --- |
| I2 | I- site double mutant | mTOR_K1753E_K1788E | Forward:<br>GAGCTTGGAGAGTGGCAGCTGAATCTACA<br>GGGCATCAATGAGAGCACAATCCCCAAAG<br>TGCTGCAGTACTACAGCGCCGCCACAGA<br>GCACGACCGCAGCTGGTACGAGGCCTGG<br>CATG<br>Reverse:<br>CAGGAAGCATCGGGCCATGAGCTTGTGCA<br>GTTCCTGCTTA |
| I3 | I- site triple mutant | mTOR_R1628E_K1655E_K1662E | Forward:<br>GAGATCGTAGAGGACTGGCAGAAAATCCT<br>TATGGTGCGGTCCCTTGTGGTCAGCCCTC<br>ATGAAGACATGAGAACCTGGCTCGAGTAT<br>GCAAGCCTGTGCGGCGAGAGTGGCAGGC<br>TGG<br>Reverse:<br>CTGGCAGCCCTGCAGTCTCTCCCACCAGA<br>T |
| I5 | I- site quintuple mutant | mTOR_R1628E_K1655E_K1662E<br>_K1706E_K1735E | Restriction and ligation of synthetic DNA fragment |
| A3 | A- site triple mutant | Rictor_R572E_R575E_R576E | Forward:<br>GAGTTTGTAGAGGAGCTACTTTATTTTTAC<br>AAGCCCAGCAGTA<br>Reverse:<br>GTGTAACTGTTCATCTTTATAGTTTCTTAG |
| A4 | A- site quadruple mutant | Rictor_R572E_R575E_R576E_Y57<br>9A | Forward:<br>GAGTTTGTAGAGGAGCTACTTGCATTTTAC<br>AAGCCCAGCAGTAAATTATAT<br>Reverse:<br>GTGTAACTGTTCATCTTTATAGTTTCTTAG |
| A5 | A- site quintuple mutant | Rictor_R572E_R575E_R576E_Y57<br>9A_L587W | Forward:<br>GAGTTTGTAGAGGAGCTACTTGCATTTTAC<br>AAGCCCAGCAGTAAATGG TATGCCAACCT<br>GGAT<br>Reverse:<br>GTGTAACTGTTCATCTTTATAGTTTCTTAG |

**Table S2.**

| | Non-uniform refinement<br><br>(EMDB-xxxx)<br>(PDB xxxx) | Focus refinement<br>on one half<br>(EMDB-xxxx)<br>(PDB xxxx) |
|---|---|---|
| **Data collection and processing** | | |
| Magnification | 59500x | 59500x |
| Voltage (kV) | 300 | 300 |
| Electron exposure (e–/Å$^2$) | ~70 | ~70 |
| Defocus range (μm) | 1.0-3.0 | 1.0-3.0 |
| Pixel size (Å) | 1.34 (1.6x binned) | 1.34 (1.6x binned) |
| Symmetry imposed | C1 | C1 |
| Initial particle images (no.) | 656’621 | 656’621 |
| Final particle images (no.) | 293’038 | 293’038 |
| Map resolution (Å) | 3.2 | 3.0 |
| FSC threshold | 0.143 | 0.143 |
| Map resolution range (Å) | 2.7-~7 | 2.7-~7 |
| **Refinement** | | |
| Initial model used (PDB code) | 5ZCS (mTOR+mLST8) | 5ZCS (mTOR+mLST8) |
| Model resolution (Å) | 3.4 | 3.2 |
| FSC threshold | 0.5 | 0.5 |
| Model resolution range (Å) | 3.2 - | 3.0 - |
| Map sharpening *B* factor (Å$^2$) | 97.47 | 69.09 |
| Model composition | | |
| Non-hydrogen atoms | 56947 | 24994 |
| Protein residues | 7437 | 3126 |
| Ligands | 8 | 4 |
| *B* factors (Å$^2$) | | |
| Protein | 122.01 | 42.44 |
| Ligand | 132.774 | 59.92 |
| R.m.s. deviations | | |
| Bond lengths (Å) | 0.004 | 0.002 |
| Bond angles (°) | 0.808 | 0.513 |
| Validation | | |
| MolProbity score | 1.57 | 1.68 |
| Clashscore | 3.83 | 6.22 |
| Poor rotamers (%) | 1.59 | 1.3 |
| Ramachandran plot | | |
| Favored (%) | 96.30 | 96.29 |
| Allowed (%) | 3.68 | 3.71 |
| Disallowed (%) | 0.03 | 0.00 |

Refinement statistics for mTORC2 half and complete complex.

**Movie S1.**

Visualization of the first and second component of the variability analysis done in Cryosparc v2, showing the continuous motion found within the particle set.

**Movie S2.**

Overview of the human mTORC2 complex, followed by a close-up view of the I-site, the binding site of InsP6 common to mTORC1 and mTORC2.

**Movie S3.**

Overview of the domain architecture of Rictor in mTORC2. ATPγS binds the A-site in the HEAT-like domain (HD) of Rictor.

**Movie S4.**

Overview of SIN1 integration into mTORC2. The SIN1 N-terminus tightly interacts with Rictor. SIN1 traverses the catalytic site cleft and winds around mLST8.
